# Cellular RNA Interacts with MAVS to Promote Antiviral Signaling

**DOI:** 10.1101/2023.09.25.559083

**Authors:** Nandan S. Gokhale, Kim Somfleth, Matthew G. Thompson, Russell K. Sam, Daphnée M. Marciniak, Lan H. Chu, Moonhee Park, Steve Dvorkin, Andrew Oberst, Stacy M. Horner, Shao-En Ong, Michael Gale, Ram Savan

**Author notes:** Corresponding authors: Nandan S. Gokhale and Ram Savan.

## Abstract

Immune signaling needs to be well-regulated to promote clearance of pathogens, while preventing aberrant inflammation. Interferons (IFNs) and antiviral genes are activated by the detection of viral RNA by RIG-I-like receptors (RLRs). Signal transduction downstream of RLRs proceeds through a multi-protein complex organized around the central adaptor protein MAVS. Recent work has shown that protein complex function can be modulated by RNA molecules providing allosteric regulation or acting as molecular guides or scaffolds. Thus, we hypothesized that RNA plays a role in organizing MAVS signaling platforms. Here, we show that MAVS, through its central intrinsically disordered domain, directly interacts with the 3′ untranslated regions of cellular mRNAs. Importantly, elimination of RNA by RNase treatment disrupts the MAVS signalosome, including newly identified regulators of RLR signaling, and inhibits phosphorylation of the transcription factor IRF3. This supports the hypothesis that RNA molecules scaffold proteins in the MAVS signalosome to induce IFNs. Together, this work uncovers a function for cellular RNA in promoting signaling through MAVS and highlights a generalizable principle of RNA regulatory control of cytoplasmic immune signaling complexes.

## INTRODUCTION

Innate immunity against RNA viruses is initiated when sentinel proteins such as those in the RIG-I-like receptor (RLR) family sense viral RNA motifs to induce type I and III interferons (IFN), pro-inflammatory cytokines, and cellular defense genes which establish an antiviral state (*1*). While a robust antiviral response is necessary to restrict viral infection, its aberrant induction can lead to inflammation and autoimmune disorders (*2*). Therefore, antiviral immune signaling needs to be tightly controlled for effective defense against viruses while restricting tissue damage. Uncovering the principles by which antiviral signaling is regulated is important for developing new therapeutic strategies against both viral infection and inflammatory disorders.

Upon sensing viral RNA in the cytosol, the RLRs RIG-I and MDA5 translocate to ER-mitochondrial and other organellar contact sites where they trigger the oligomerization of the adaptor protein MAVS (*3–6*). Oligomerized MAVS acts as a platform to recruit multiple proteins, which together organize into a higher order signaling complex (termed the MAVS signalosome). At the MAVS signalosome, the kinases TBK1, IKK-ε, and the IκB complex phosphorylate and activate the transcription factors IRF3 and NF-κB (*7–12*), which then induce IFN and an antiviral gene program. The MAVS signalosome requires a series of protein-protein interactions and post-translational modifications (PTMs) at specific subcellular sites for efficient signal transduction (*1*, *13*). However, whether non-viral RNA molecules directly regulate antiviral immune signaling at the MAVS signalosome remains unknown.

RNA interacts with proteins in a sequence- and structure-dependent fashion; this ability is a key factor that drives the versatile functions of RNA, which extend well beyond its role as an intermediate between DNA and protein. RNA can recruit proteins and promote their interactions within molecular complexes. This is exemplified by ribosomal RNAs, which serve as a scaffold for dozens of interacting proteins to build the ribosome (*14*). Long non-coding RNAs and the 3′ untranslated regions (UTRs) of mRNAs can similarly modulate protein and protein complex functions by serving as molecular guides or scaffolds (*15–19*). Further, RNA binding can lead to conformational or functional changes in proteins, thereby allosterically influencing their function (*20–23*). The number of proteins that are known to interact with RNA has greatly increased through recent proteomics-based studies, and a large proportion of these newly identified RNA-binding proteins (RBPs) do not contain any canonical RNA-binding domains (*24*, *25*). This highlights the possibility of widespread RNA-mediated regulation of macromolecular protein complexes and their function. Therefore, we hypothesized that RNA could interact with proteins in the RLR pathway to modulate signaling.

In this study, we found that the central disordered domain of MAVS, the adaptor protein of RLR signaling, is associated with the 3′ UTRs of cellular mRNAs. RNA binding to MAVS promotes downstream signaling by altering the interaction of this adaptor with several new proteins found to be part of the MAVS signalosome. Together, this work highlights a previously unappreciated role for RNA-dependent regulation of the MAVS signalosome affecting antiviral immunity, which could be broadly applicable to other cytosolic immune signaling complexes.

## RESULTS

### Cellular RNA promotes IRF3 phosphorylation by the MAVS signalosome

As higher-order protein complexes can be scaffolded by RNA, we hypothesized that RNA may regulate MAVS signalosome activity. The phosphorylation of the transcription factor IRF3, which is required to induce IFNs, is a key outcome of the activation of the RLR pathway through the MAVS signalosome (*11*, *12*, *10*, *26*). To determine whether the MAVS signalosome is regulated by RNA, we tested its ability to phosphorylate IRF3 *in vitro* with RNase treatment (*27*). Here, treated mitochondrial extracts from unstimulated or stimulated 293T IRF3^KO^ cells are incubated with cytosolic extract from unstimulated 293T^WT^ cells. 293T^WT^ cytosolic extract is the only source of IRF3 in these reactions, allowing for the discrimination of *in vitro* phosphorylated IRF3 as a measure of MAVS signalosome activity (Fig. 1A).

**Fig. 1:**
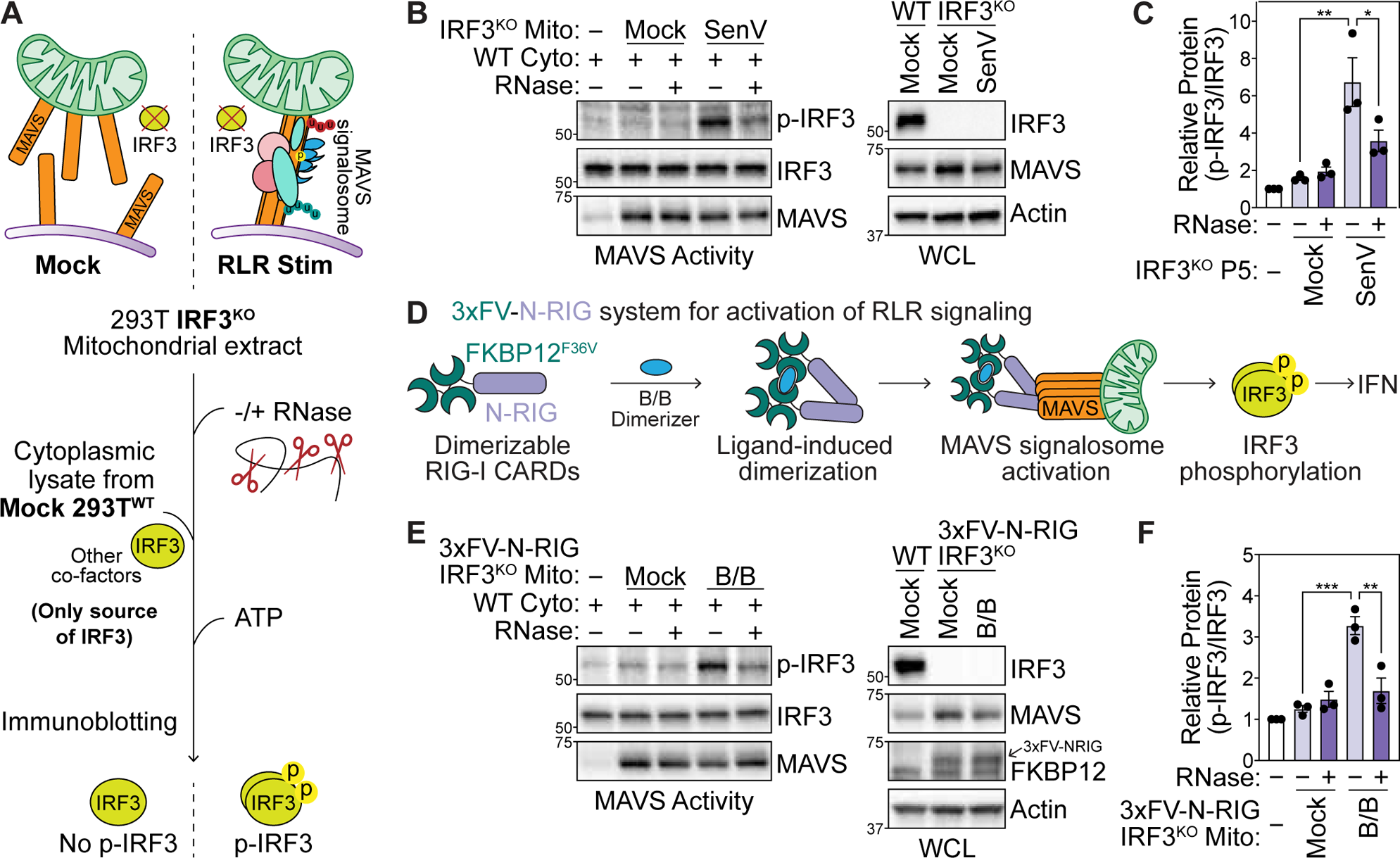
Cellular RNA promotes IRF3 phosphorylation at the MAVS signalosome. **(A)** MAVS activity assay for *in vitro* phosphorylation of IRF3. Crude mitochondrial (Mito) extracts from 293T IRF3^KO^ cells are incubated with cytoplasmic extracts (Cyto) from unstimulated 293T^WT^ cells in the presence of ATP. IRF3 phosphorylation is analyzed by immunoblot. **(B)** MAVS activity assay to analyze *in vitro* IRF3 phosphorylation by mitochondrial extracts −/+ RNase treatment from mock or SenV-infected (100 HAU/mL, 16 hpi) 293T IRF3^KO^ cells. WCL = whole cell lysates. **(C)** Quantification of p-IRF3 (S386) relative to IRF3 from experiments in (B). **(D)** Outline of 3xFV-N-RIG system that activates the MAVS signalosome in the absence of viral RNA. Tandem FKBP12^(F36V)^ dimerization domains are fused to the N-terminal CARDs of RIG-I. Treatment with the small molecule B/B multimerizes 3xFV-N-RIG to activate downstream signaling. **(E)** MAVS activity assay to analyze *in vitro* IRF3 phosphorylation by mitochondrial extracts −/+ RNase treatment from mock or B/B-treated (10 nM, 3 hpt) 293T IRF3^KO^ cells stably expressing 3xFV-N-RIG. **(F)** Quantification of p-IRF3 relative to IRF3 (S386) from experiments in (E). Data in (B) and (E) are representative of 3 biological replicates. Values in (C) and (F) are mean ± SEM of 3 biological replicates. * p ≤ 0.05, ** p ≤ 0.01, *** p ≤ 0.001 by one-way ANOVA with Tukey’s multiple comparison test.

As expected, mitochondrial extracts from Sendai (SenV; a potent activator of RIG-I) virus-infected 293T IRF3^KO^ cells, but not from uninfected cells, robustly phosphorylated IRF3 at serine 386. However, IRF3 phosphorylation by mitochondrial extracts from SenV-infected cells was inhibited by RNase treatment, which indicates that RNA promotes MAVS signalosome activity (Fig. 1B-C). For these experiments, we used a cocktail of RNase A and RNase III, which cleave single-stranded and double-stranded RNA, respectively. To further investigate which form of RNA promoted *in vitro* IRF3 phosphorylation by the MAVS signalosome, we compared this cocktail of RNases with treatment with RNase A or RNase III alone, and with DNase I treatment (Fig. S1A-B). RNase A treatment inhibited IRF3 phosphorylation to the same level as RNase A+III treatment. Conversely, neither RNase III alone nor DNase I treatment inhibited IRF3 phosphorylation. These data suggest that single-stranded RNAs or RNA regions are responsible for promoting MAVS signalosome function.

While SenV RNA within ribonucleoprotein complexes inside cells is resistant to RNase treatment, viral RNA complexed with RIG-I may still contribute to the observed RNA-modulated phosphorylation of IRF3 by the MAVS signalosome (*28*). To generate a system for measuring MAVS signalosome activity independently of viral RNA, we fused the N-terminal caspase activation and recruitment domains (CARD) of RIG-I to FKBP12^F36V^ dimerizing domains (3xFV-N-RIG). Here, the small non-RNA molecule B/B multimerizes 3xFV-N-RIG, which is sufficient to activate the MAVS signalosome and propagate downstream signaling (Fig. 1D) (*29*, *30*). Indeed, B/B treatment rapidly induces *IFNB1* in 293T cells stably expressing 3xFV-N-RIG, but not in 293T^WT^ cells (Fig. S1C). To test whether RNase treatment inhibits IRF3 phosphorylation in the absence of an activating non-self RNA ligand, we performed the MAVS activity assay as above in 293T IRF3^KO^ that stably express 3xFV-N-RIG. While mitochondrial extracts from B/B-treated cells phosphorylated IRF3 *in vitro*, RNase treatment of these extracts abolished IRF3 phosphorylation, matching our observations during SenV infection (Fig. 1E-F). Together, these data confirm that cellular, and not viral, RNA promotes MAVS signalosome activity at the level of IRF3 phosphorylation.

### The MAVS signalosome is associated with cellular RNA

Given that cellular RNA regulates antiviral signaling through MAVS, we hypothesized that MAVS and its signalosome are associated with RNA. RNA association with proteins and protein complexes can be determined by a ribonuclease (RNase)-dependent shift in their migration through a sucrose gradient (*31*). Due to the higher mass conferred by bound RNA, proteins associated with RNA are found in higher sucrose (denser) fractions in the absence of RNase and migrate to lower sucrose (less dense) fractions in the presence of RNase (Fig. 2A). As MAVS is localized on mitochondria and components of the endomembrane system, we first isolated crude mitochondrial fractions to enrich for MAVS and its interacting partners (Fig. S2A-B). To test whether MAVS exhibits an RNase-dependent shift, we treated mitochondrial lysates from uninfected and SenV-infected 293T cells with RNases, the separated these lysates by ultracentrifugation through sucrose gradients and measured the migration of MAVS signalosome proteins by immunoblotting of collected fractions. As expected, SenV infection caused MAVS to migrate to heavy fraction 10 due to its oligomerization and the formation of the MAVS signalosome (Fig. 2B-C). MAVS in heavy, and not light, fractions can transduce signal to IRF3 (*6*, *32*). RNase treatment reduced MAVS migration to heavy fraction 10, just as it did for the canonical RBP HuR (Fig. 2B-C). Additionally, RNase treatment reduced the co-sedimentation of key components of the MAVS signalosome, including TRAF2 and TRAF3, at heavy fractions with MAVS during SenV infection, indicating that the mass of the entire complex is reduced by RNase treatment, and that one or more of the proteins in the MAVS signalosome are associated with RNA. Given that RIG-I interacts with SenV RNA, the RNase-dependent shift for this sensor is expected in infected cells (*33*). Migration of CANX, a protein not known to interact with RNA, was unaffected by RNase treatment in these experiments.

**Fig. 2:**
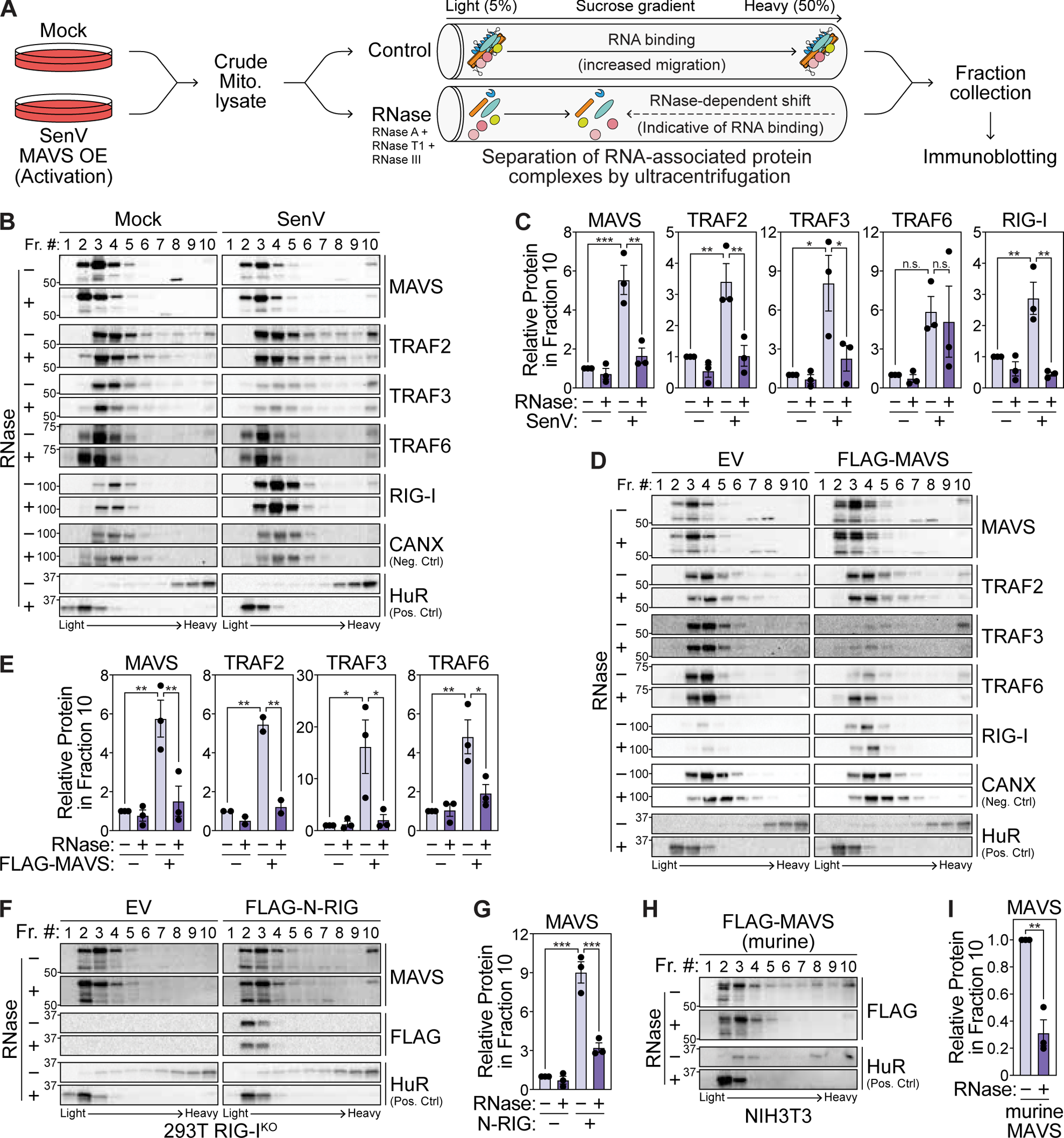
The MAVS signalosome is associated with cellular RNA. **(A)** RNase-dependent shift assay to identify RNA-associated proteins. Mitochondrial lysates treated with RNase inhibitor (−RNase) or an RNase cocktail (+ RNase; RNase A, I, T1, III) are separated on a sucrose gradient by ultracentrifugation. RNA-associated proteins have reduced migration to heavy fractions in the presence of RNase. **(B)** Immunoblot analysis of fractions collected after sucrose gradient ultracentrifugation of mitochondrial lysates −/+ RNase treatment from mock- and SenV-infected (100 HAU/mL, 14 hpi) 293T cells. **(C)** Quantification of the indicated protein in fraction 10 relative to that in fraction 3 from experiments in (B). **(D)** Immunoblot analysis of fractions collected after sucrose gradient ultracentrifugation of mitochondrial lysates −/+ RNase treatment from 293T cells transfected with empty vector (EV) and FLAG-tagged MAVS (24 hpt). **(E)** Quantification of the indicated protein in fraction 10 relative to that in fraction 3 from experiments in (D). **(F)** Immunoblot analysis of fractions collected after sucrose gradient ultracentrifugation of mitochondrial lysates −/+ RNase treatment from 293T RIG-I^KO^ cells transfected with empty vector and FLAG-tagged N-RIG. **(G)** Quantification of MAVS in fraction 10 relative to that in fraction 3 from experiments in (F). **(H)** Immunoblot analysis of fractions collected after sucrose gradient ultracentrifugation of mitochondrial lysates −/+ RNase treatment from murine NIH3T3 cells transfected with FLAG-tagged murine MAVS. **(I)** Quantification of FLAG-tagged murine MAVS in fraction 10 relative to that in fraction 3 from experiments in (H). Data in (B), (D), (F), and (H) are representative of 3 biological replicates. Values in (C), (E), (G), and (I) are mean ± SEM of 3 biological replicates. * p ≤ 0.05, ** p ≤ 0.01, *** p ≤ 0.001 by one-way ANOVA with Tukey’s multiple comparison test ((C), (E), and (G)) or unpaired t test (I). n.s. = not significant.

MAVS overexpression is sufficient to drive downstream signaling in the absence of viral infection. MAVS, TRAF2, TRAF3, and TRAF6 also demonstrated an RNase-dependent shift following overexpression of FLAG-tagged MAVS (Fig. 2D-E). However, RIG-I did not co-sediment with MAVS in the heavy fraction during MAVS overexpression, likely due to the absence of interacting viral RNA, suggesting that viral RNA bound to RIG-I is not responsible for the RNase-dependent shift of MAVS signalosome components. Indeed, MAVS also has an RNase-dependent shift in 293T RIG-I^KO^ cells transfected with the N-terminal domains of RIG-I (N-RIG) which activates signaling but does not interact with RNA (Fig. 2F-G and S2C) (*30*). FLAG-tagged murine MAVS overexpressed in mouse NIH3T3 cells also exhibited an RNase-dependent shift, demonstrating that MAVS-RNA association is conserved between human and murine cells (Fig. 2H-I). Together, these data indicate that higher order MAVS complexes are abolished by RNase treatment, indicating that the MAVS signalosome is associated with cellular RNA.

MAVS oligomerization also increases the migration of this protein to higher sucrose fractions (*6*, *32*). To ensure that the RNase-dependent shift of the MAVS signalosome was not due to a resolution of MAVS oligomers, we tracked protein oligomers using semi-denaturing detergent agarose gel electrophoresis (SDD-AGE) on mitochondrial lysates isolated from uninfected or SenV-infected 293T cells with or without RNase treatment. SenV infection led to the formation of MAVS oligomers, which were not resolved by RNase treatment. On the other hand, β-mercaptoethanol (β-ME) resolved MAVS oligomers as expected (Fig. S2D). Similarly, oligomers formed by MAVS overexpression were resolved by β-ME but were refractory to RNase treatment (Fig. S2E). Therefore, the observed RNase-dependent shift of the MAVS signalosome is due to RNA association, and not due to MAVS de-oligomerization.

### MAVS interacts with cellular RNA through its conserved central disordered domain

Infrared-dye crosslinking and immunoprecipitation (irCLIP) allows for the detection of RNA-protein complexes. Following stringent immunoprecipitation of the target protein, an infrared-dye fluorescent oligonucleotide is ligated to UV-crosslinked RNA fragments and RNA-protein complexes are visualized by SDS-PAGE (Fig. 3A) (*34*). To test whether MAVS directly interacts with RNA, we performed irCLIP on FLAG-tagged MAVS expressed in 293T MAVS^KO^ cells and detected crosslinking-dependent MAVS-RNA complexes (Fig. 3B-C and S2A). irCLIP of endogenous MAVS in uninfected and SenV-infected 293T cells revealed that MAVS interacts with RNA in both uninfected and infected cells (Fig. 3D-E). Further, overexpressed FLAG-tagged murine MAVS also interacts with RNA in mouse NIH3T3 cells, suggesting that MAVS-RNA interaction is conserved between humans and mice (Fig. 3F-G).

**Fig. 3:**
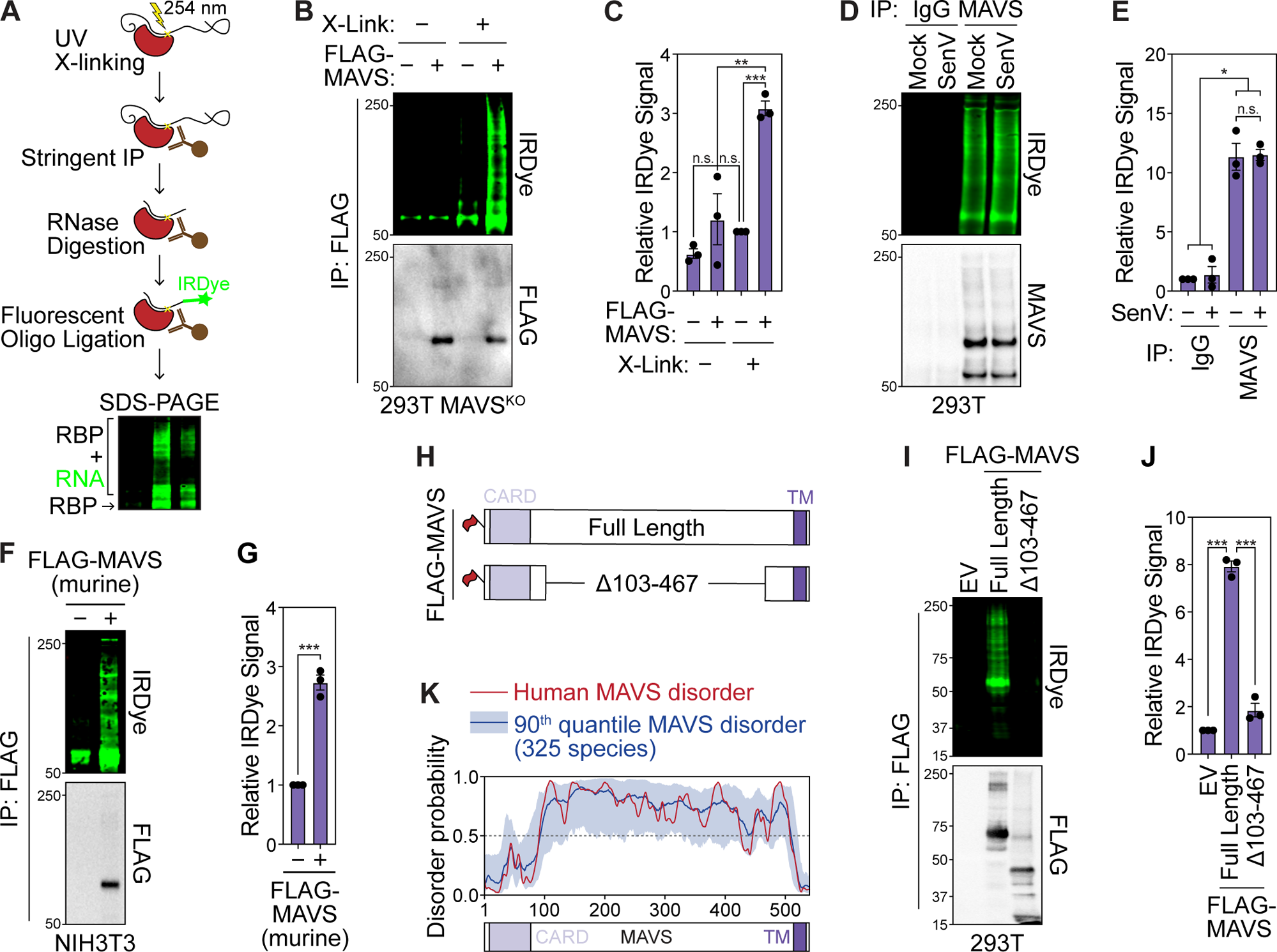
MAVS interacts with RNA through its central intrinsically disordered region. **(A)** Infrared-dye crosslinking and immunoprecipitation (irCLIP) strategy to visualize RNA-protein complexes. The protein of interest is stringently immunoprecipitated and UV-crosslinked RNA is digested to fragments with RNase A. Following ligation of an IRDye-800 conjugated oligonucleotide, complexes are resolved by SDS-PAGE. RNA-protein complexes are detected by IRDye-800 fluorescence and immunoprecipitation is validated by immunoblot analysis. **(B)** irCLIP −/+ crosslinking of FLAG-tagged MAVS expressed in 293T MAVS^KO^ cells (24 hpt). EV = empty vector. **(C)** Quantification of IRDye signal in experiments in (B). **(D)** irCLIP of endogenous MAVS from mock- and SenV-infected (100 HAU/mL, 16 hpi) 293T cells. **(E)** Quantification of IRDye signal in experiments in (D). **(F)** irCLIP of FLAG-tagged murine MAVS expressed in murine NIH3T3 cells (24 hpt). **(G)** Quantification of IRDye signal in experiments in (F). **(H)** Schematic of FLAG-tagged full length MAVS and MAVSΔ103-467 used in (I). **(I)** irCLIP of the indicated FLAG-tagged MAVS constructs expressed in 293T MAVS^KO^ cells (24 hpt). **(J)** Quantification of IRDye signal in experiments in (I). **(K)** Prediction of disorder in human MAVS (red) and in 325 mammalian species (blue) using IUPred3. Data in (B), Data in (D), (F), and (I) are representative of 3 biological replicates. Values in (C), (E), and (G), and (J) are mean ± SEM of 3 biological replicates. * p ≤ 0.05, ** p ≤ 0.01, *** p ≤ 0.001 by one-way ANOVA with Tukey’s multiple comparison test. n.s. = not significant.

To identify the domains of MAVS responsible for interacting with RNA, we performed irCLIP in 293T MAVS^KO^ cells by expressing FLAG-tagged human MAVS constructs with deletions in the caspase activation and recruitment domain (CARD), transmembrane domain (TM), or the central region between amino acids 103 and 467 (Fig. S3B). The CARD is required for MAVS oligomerization and activation, while the TM domain mediates MAVS localization of mitochondrial and other cellular membranes (*11*, *13*). The central region of MAVS between amino acids 103-467 harbors binding sites for many MAVS-interacting proteins, including TRAF2, TRAF3, TRAF6, and IRF3 (*7*, *9*, *13*, *35*). As expected, overexpression of these mutants in 293T MAVS^KO^ cells did not induce signaling (Fig. S3C-D). Deletion of the CARD, TM, or both CARD and TM did not reduce MAVS-RNA interaction as determined by irCLIP (Fig. S3E-F). However, deletion of amino acids 103-467 of MAVS eliminated the ability of this protein to bind RNA, which indicates that MAVS interacts with RNA through this region (Fig. 3H-J and S3E-F).

This central region of MAVS is predicted to harbor extensive intrinsic disorder by two different prediction softwares (IUPred3 and FuzPred) as well as AlphaFold, and is enriched for disorder-promoting residues like serine, proline, and glycine (Fig. 3K and Fig. S3G-I) (*36–38*). Further, this disorder across the central region of MAVS was predicted to be conserved across 325 species (Fig. 3K). Intrinsically disordered domains are known to promote protein-RNA interactions (*25*, *39–42*). Together, these data indicate that MAVS directly interacts with RNA through its conserved, central disordered domain.

### MAVS interacts with the 3′ UTR of mRNAs

Traditional UV-crosslinking and immunoprecipitation-based techniques to identify bound RNAs are challenging for proteins that lack canonical RNA-binding domains, such as MAVS (*43*, *44*). Therefore, we used targeted RNA editing mediated by APOBEC1 to identify MAVS-associated RNAs. Targeted APOBEC1-mediated editing can robustly identify the RNA interactomes of RBPs without relying on crosslinking or immunoprecipitation. Here, APOBEC1 fused to the protein of interest catalyzes C-to-U transitions in RNAs that interact with this protein, which can be detected as C-to-T mutations by RNA sequencing (Fig. 4A) (*45*, *46*).

**Fig. 4:**
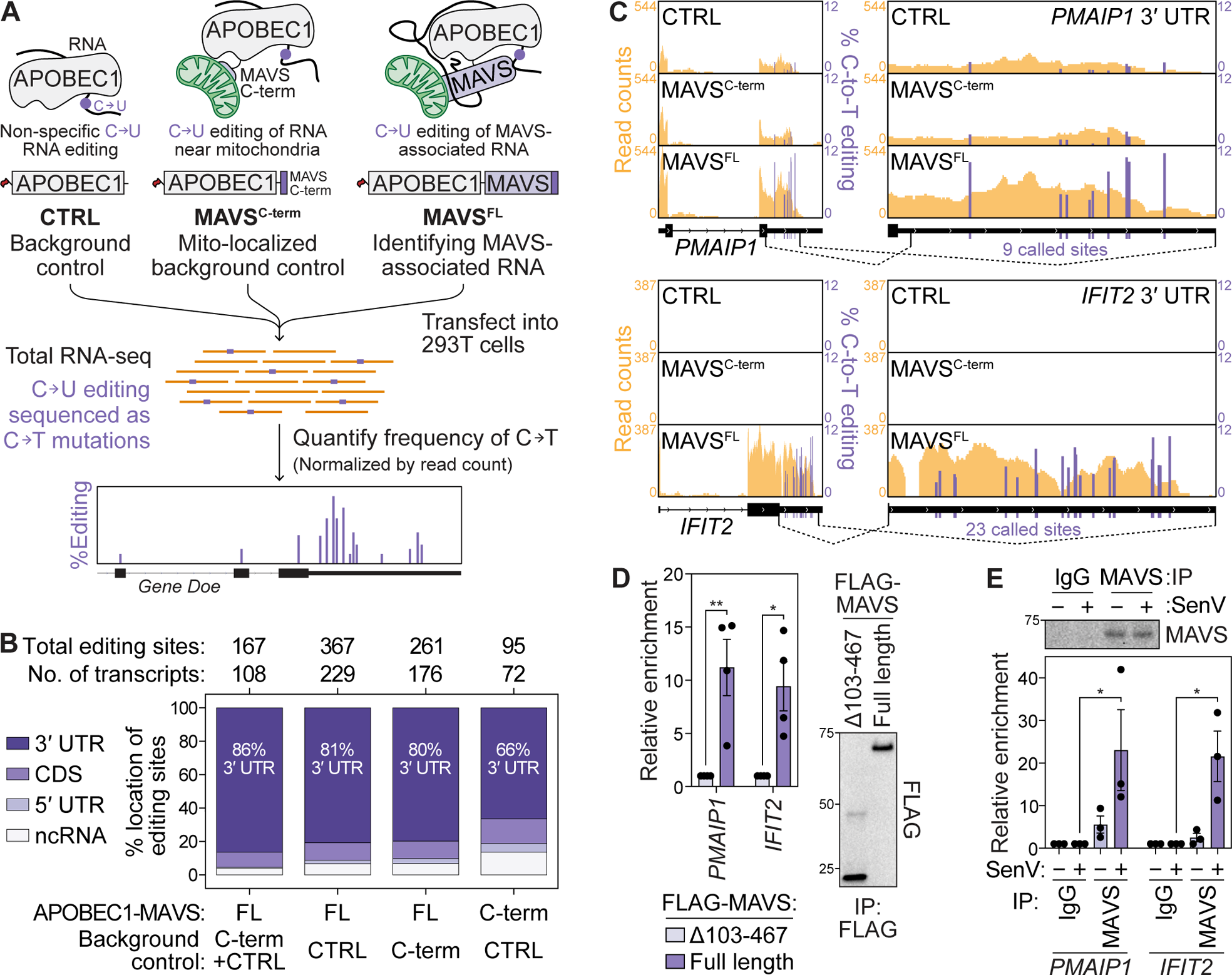
MAVS interacts with the 3′ UTRs of mRNAs. **(A)** Targeted APOBEC1-mediated editing approach used to profile MAVS-interacting RNAs. C-to-T edits in sequenced cDNA enhanced with APOBEC1-MAVS^FL^ overexpression relative to non-specific (APOBEC1 alone; CTRL) and mitochondria-localized (APOBEC1-MAVS^C-term^) background controls identifies MAVS-associated RNAs. **(B)** Summary of C-to-T editing sites and edited transcripts identified when comparing the indicated APOBEC1-MAVS constructs with background controls across biological triplicates. **(C)** Percent C-to-T editing (purple) at called sites in *PMAIP1* and *IFIT2* 3′UTRs (mean of 3 replicates). Read counts from one representative experiment are shown in yellow. **(D)** Left: CLIP-RT-qPCR analysis of normalized enrichment relative to input of *PMAIP1* and *IFIT2* mRNA by immunoprecipitation of the indicated FLAG-tagged MAVS constructs in transfected 293T MAVS^KO^ cells (24 hpt). Values are mean ± SEM of 4 biological replicates. * p ≤ 0.05, ** p ≤ 0.01, by unpaired t test. Right: Representative immunoblot of immunoprecipitated fractions. **(E)** Bottom: CLIP-RT-qPCR analysis of normalized enrichment relative to input of *PMAIP1* and *IFIT2* mRNA by immunoprecipitation of endogenous MAVS from mock- and SenV-infected (100 HAU/mL, 16 hpt) 293T cells. IgG was used as background control. Values are mean ± SEM of 3 biological replicates. * p ≤ 0.05 by one-way ANOVA with Tukey’s multiple comparison test. Top: Representative immunoblot of immunoprecipitated fractions.

To profile MAVS-associated RNAs, we fused FLAG-tagged APOBEC1 to the N-terminus of full-length MAVS (APOBEC1-MAVS^FL^). We used FLAG-APOBEC1 as a background control (CTRL), and FLAG-APOBEC1 fused to the C-terminal transmembrane helix of MAVS (MAVS^C-term^) as a mitochondrially-localized background control to account for off-target editing within the subcellular vicinity of MAVS (Fig. 4A). Transfection of APOBEC1-MAVS^FL^ into 293T cells induced *IFNB1*, indicating that this construct can form the MAVS signalosome, while APOBEC1 and APOBEC1-MAVS^C-term^ did not induce *IFNB1* (Fig. S4A-B). Further, while APOBEC1 was localized primarily to the cytoplasm, both APOBEC1-MAVS^FL^ and APOBEC1-MAVS^C-term^ had a similar membrane-associated pattern of localization (Fig. S4C).

RNA-sequencing data from 293T cells transfected with these constructs were analyzed using the Bullseye pipeline to identify RNAs with increased C-to-T editing with APOBEC1-MAVS^FL^ as compared to controls (Supplementary Data S1) (*46*, *47*). We identified enhanced C-to-T editing at a set of 167 sites in 107 transcripts when comparing APOBEC1-MAVS^FL^ to both controls together, and further editing sites emerged when comparing APOBEC1-MAVS^FL^ to each control individually (Fig. 4B and Table S1). Importantly, the low overlap of edited transcripts between APOBEC1-MAVS^FL^ and APOBEC1-MAVS^C-term^ when compared to APOBEC1, and the higher number of transcripts modified by APOBEC1-MAVS^FL^, indicates that APOBEC1-MAVS^FL^ specifically edits MAVS-associated transcripts (Fig. 4B and Fig. S4D). Most editing sites (86%) were identified in the 3′ untranslated regions (UTRs) of mRNAs.

Given that MAVS interacts with over 100 mRNAs, we do not hypothesize that any one transcript is a primary driver of RNA-regulated modulation of MAVS function. However, the 3′ UTRs of *PMAIP1* and *IFIT2*, which contain 9 and 23 editing sites respectively, suggest that these UTRs are highly associated with MAVS (Fig. 4C and Fig. S4E). The rate of editing of called sites in *PMAIP1* by APOBEC1-MAVS^FL^ as determined by Sanger sequencing was correlated with that determined by RNA-seq. (Fig. S4F-G). *PMAIP1* and *IFIT2* binding to MAVS was further validated by CLIP-RT-qPCR. Both transcripts were enriched with full length FLAG-MAVS as compared to FLAG-MAVS Δ103-467, which does not bind RNA, when these constructs were overexpressed in 293T MAVS^KO^ cells (Fig. 4D). Both *PMAIP1* and *IFIT2* are induced by IFNs (Fig. 4C) (*48*). This may indicate a feedback mechanism wherein some newly transcribed IFN-inducible mRNAs bind to MAVS. Indeed CLIP-RT-qPCR, when normalized to input RNA levels, indicates that the interaction of endogenous MAVS with *PMAIP1* and *IFIT2* mRNAs is augmented by SenV infection (Fig. 4F). Taken together, these data suggest that MAVS interacts primarily with the 3′ UTRs of cellular RNAs, and that specific MAVS-RNA interactions may be altered during active signaling.

### RNA modulates functional MAVS protein-protein interactions

As MAVS interacts with RNA and RNase treatment inhibits *in vitro* IRF3 phosphorylation by the MAVS signalosome, we hypothesized that RNA alters the ability of MAVS to interact with proteins required for proper signaling function. To identify proteins that differentially interact with MAVS in an RNA-dependent manner, we immunoprecipitated (IP) FLAG-MAVS from mitochondrial lysates from transfected 293T cells with or without RNase treatment and analyzed immunoprecipitated MAVS complexes by mass spectrometry (MS) (Fig. 5A). Mitochondrial lysates from empty vector-transfected cells without RNase treatment were used as background control.

**Fig. 5:**
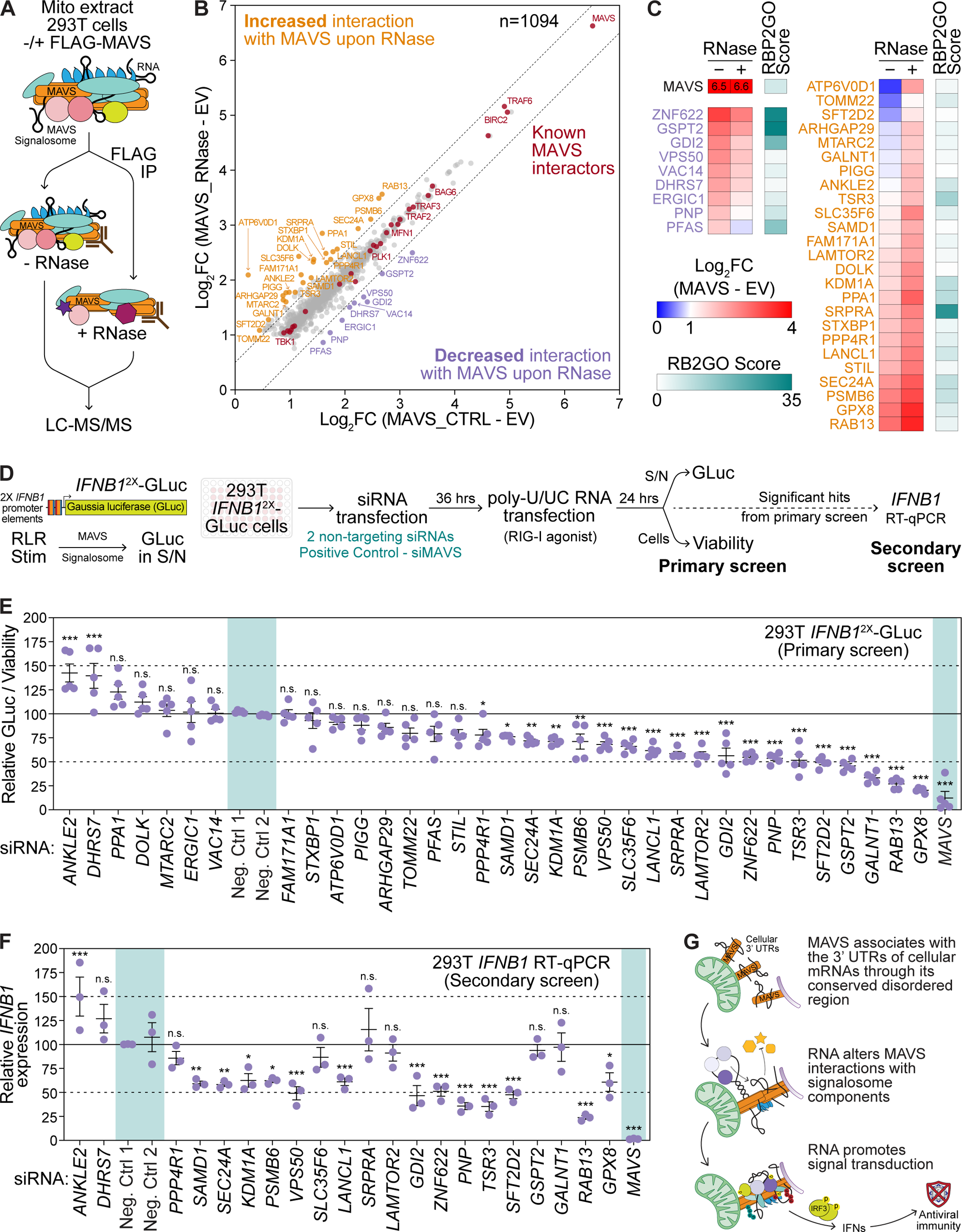
RNA alters MAVS protein-protein interactions. **(A)** Immunoprecipitation and mass spectrometry strategy to identify RNA-dependent MAVS interactors. **(B)** Scatterplot of 1094 proteins significantly enriched with MAVS (Log_2_FC ≥ 2, p ≤ 0.05, found in < 30% of CRAPome datasets) −/+ RNase over empty vector control across 5 biological replicates. Dashed lines delineate |Log_2_FC| = 0.5 from the diagonal. Known MAVS-interacting proteins are in red, those with decreased MAVS-interaction upon RNase treatment are in purple, and those with increased MAVS-interaction upon RNase treatment are in ochre. **(C)** Heatmap of the enrichment of RNA-dependent MAVS-interactors over empty vector control −/+ RNase (blue-red gradient), and the RBP2GO score, a measure of empirically determined propensity to interact with RNA, for each protein (teal). **(D)** Schematic of targeted siRNA screen for RNA-dependent MAVS interactors. **(E)** Relative *Gaussia* luciferase (GLuc) activity normalized to viability following poly-U/UC RNA transfection (50 ng, 24 hpt) in 293T *IFNB1*^2X^-GLuc reporter cells upon depletion of RNA-dependent MAVS interactors by siRNA treatment (36 hours). Controls are highlighted in teal. **(F)** Normalized *IFNB1* mRNA expression relative to *HPRT1* following poly-U/UC RNA transfection (50 ng, 24 hpt) in 293T cells upon depletion of RNA-dependent MAVS interactors by siRNA treatment (36 hours), determined by RT-qPCR. Controls are highlighted in teal. **(G)** Proposed model for RNA regulation of MAVS signalosome function. Values are the mean ± SEM of 5 (E) or 3 (F) biological replicates. * p ≤ 0.05, ** p ≤ 0.01, *** p ≤ 0.001 by one-way ANOVA with Tukey’s multiple comparison test. n.s. = not significant.

We identified a total of 3200 proteins across 5 biological replicates, of which 1094 proteins were significantly enriched (≥ 2-fold, p ≤ 0.05, detected in < 30% of datasets in the CRAPome database) with FLAG-MAVS −/+ RNase treatment over empty vector control (Fig. 5B, Fig. S5A, and Supplementary Data S2). These include proteins known to interact with MAVS and regulate its function, such as core signalosome components like TRAF2, TRAF3, TRAF6, and TBK1, highlighting the specificity of this IP-MS strategy (*7*, *9*, *13*, *49*). Functional network analysis with STRING revealed that many MAVS interacting proteins fell into discrete protein complexes or functional categories (Fig. S5B) (*50*). Gene Set Enrichment Analysis further revealed that these proteins were enriched in pathways related to immune responses, signaling cascades, metabolism, organellar organization, transport, ER stress, and post-translational modification (Fig. S5C) (*51*, *52*).

While most identified MAVS-interacting proteins were similarly enriched with or without RNase treatment, 9 showed decreased and 25 showed increased (Log_2_FC > |0.5|) interaction with MAVS following RNase treatment (Fig. 5B-C). Proteins demonstrating reduced MAVS interaction with RNase treatment had a higher RBP2GO score on average, indicating that these proteins are more likely to interact with RNA (Fig. 5C). The RBP2GO score is calculated based on the number of times a factor is identified in proteomic RBP discovery datasets, and on its interaction networks with other RBPs (*24*). On the other hand, proteins with increased MAVS interaction upon RNase have a lower RBP2GO score on average.

To identify whether these RNA-altered MAVS interactors regulate RLR signaling, we performed a targeted siRNA screen in 293T cells stably expressing a *IFNB1*^2X^-*Gaussia* luciferase (GLuc) reporter (Fig. 5D). Here, secreted GLuc, kept basally low by a destabilizing PEST domain, is induced by two tandem enhanceosome elements from the *IFNB1* promoter (Fig. S6A) (*53*, *54*). Clonal 293T *IFNB1*^2X^-GLuc cells demonstrate dose-dependent GLuc induction in response to an RLR ligand such as transfected poly-U/UC, an *in vitro*-transcribed, immunogenic RNA fragment from the hepatitis C virus genome that potently activates RIG-I (Fig. S6B) (*55*). Further, GLuc induction upon poly-U/UC RNA transfection is inhibited by depletion of MAVS or IRF3, indicating that this reporter is sensitive to perturbation of the RLR pathway (Fig. S6C).

Depletion of 21/34 newly identified RNA-modulated MAVS-interacting proteins significantly affected *IFNB1* promoter-driven GLuc production in response to poly-U/UC RNA, with 2 increasing and 18 decreasing GLuc production (Fig. 5E). To account for possible effects on GLuc transcription, translation, or secretion, we then performed a secondary screen on hits that altered *IFNB1*^2X^-GLuc production, testing for the induction of *IFNB1* mRNA by poly-U/UC in 293T cells (Fig. 5F). From these screens, we determined that depletion of 4/9 proteins with reduced interaction with MAVS upon RNase treatment (PNP, ZNF622, GDI2, and VPS50) decreased both GLuc and *IFNB1* induction, indicating that these factors promote signaling. On the other hand, 9/25 proteins with increased MAVS interaction upon RNase treatment (GPX8, RAB13, SFT2D2, TSR3, LANCL1, PSMB6, KDM1A, SEC24A, SAMD1) also promote signaling. ANKLE2, whose interaction with MAVS is increased by RNase treatment, inhibits RLR signaling. Together, these data indicate that proteins that associate with MAVS in an RNA-regulated manner can promote or inhibit antiviral signaling through the RLR pathway. To our knowledge, none of these factors has yet been found to play a role in RLR signaling and instead represent novel regulatory components of the MAVS signalosome.

## DISCUSSION

Here we uncover a new role for cellular RNA in promoting RLR signaling through interaction with the adaptor protein MAVS. We demonstrate that the MAVS signalosome is associated with RNA, and that MAVS directly binds to the 3′ UTRs of cellular mRNAs through its central disordered domain. RNA binding alters the ability of MAVS to interact with several newly identified protein interactors, thereby modulating signaling (Fig. 5G). Therefore, IFN induction through the RLR pathway is initially triggered by pathogenic self- or non-self RNA, but also augmented by cellular RNA binding to MAVS.

Although MAVS does not harbor a canonical RNA-binding domain, we found that MAVS interacts with the UTRs of cellular mRNAs through its central disordered region. Disordered domains can directly interact with RNA; a large proportion of proteins newly recognized to interact with RNA do not contain canonical RNA-binding domains but do encode disordered regions. Such interactions are common as roughly half of all peptides that can be crosslinked to RNA map to disordered regions (*25*, *56*, *57*). Typically, disordered RNA-binding domains are enriched in positively charged arginine- or lysine-rich basic patches, or arginine/glycine and arginine/serine repeats which encourage interaction with the negatively charged phosphate backbone of RNA (*41*, *56*). Intriguingly, the MAVS disordered region, which is enriched for disorder-promoting amino acids including glycine, serine, and proline, is depleted for positively charged residues. This suggests a new mode of non-canonical interaction between the MAVS disordered region and RNA, which will be evaluated in future experiments.

Previous studies have shown that protein-protein interactions form the MAVS signalosome, initially nucleated by CARD-mediated oligomerization of MAVS itself. These studies have revealed critical effectors such as TRAF2, TRAF3, TRAF6, TBK1, and ATP13A1, all of which we also identified in our proteomics approach, which are required for the activation of IFNs (*7–9*, *58*, *59*). Our study expands these functional interactors, uncovering new proteins that associate with MAVS in an RNA-modulated manner to influence IFN induction. Future work will determine the precise mechanisms by which RNA mediates these interactions, and how these factors modulate signaling.

We propose two potential mechanisms for how RNA could affect these protein-protein interactions with MAVS. First, cellular RNA could directly promote or inhibit interactions with these newly discovered regulators by simultaneously interacting with MAVS and RNA-binding factors that promote signaling such as ZNF622 and GDI2, thereby bringing these proteins into proximity for functional interaction. Similarly, RNA could sterically prevent interactions between MAVS and factors that inhibit signaling such as ANKLE2 or change the temporal order of signalosome coalescence required for efficient signaling. Second, RNA interaction may change the conformation of the MAVS disordered region itself, thereby promoting or inhibiting protein binding. Disordered regions are flexible and can undergo local disorder-to-order transitions upon ligand binding, which can stabilize and functionalize neighboring segments. For example, RNA binding changes the conformational state of the disordered N-terminal domain of MYC, thereby potentiating the formation of functional MYC complexes (*23*). Therefore, RNA-dependent conformational changes in MAVS may stabilize or destabilize short linear motifs within the conserved disordered region to modulate interaction with other signalosome components. Indeed, specific binding motifs for TRAF proteins and IRF3 within the MAVS disordered region are already known to be exposed and functionalized by CARD-mediated MAVS oligomerization (*6*, *35*). Except for a few motifs in the disordered region, to which TRAF proteins and IRF3 bind, and previously discovered sites of MAVS post-translational modification, the functional significance of the evolutionarily conserved disordered region in MAVS has been unclear. Our study reveals that RNA binding to this disordered region adds an important regulatory layer to MAVS function.

While post-transcriptional regulation of RNA has been extensively studied in the context of immune responses, the scaffolding functions of RNA in immune signaling complexes are limited. Here, we describe a role for RNA in modulating the MAVS signalosome and its function and have identified novel RNA-dependent protein regulators that affect downstream IFN activation. Further studies will be required to interrogate precisely how these effectors modulate MAVS activity. Interestingly, we find both positive and negative regulators of MAVS activity, suggesting that activation of this pathway is tightly regulated at the adaptor level. The conservation of disorder in MAVS indicates that RNA-dependent functions are conserved across humans and mice. Therefore, the concept that RNA could modulate the activation of higher-order protein complexes, which are a feature of innate immune signaling (*60*), may broadly apply to immune signaling adaptors.

## ACKNOWLEDGEMENTS

We would like to thank Dr. Nicholas Heaton, Dr. Jennifer Hyde, Dr. Kate Meyer, Dr. Yasemin Sancak, Dr. Christine Vazquez, Dr. Kevin Labagnara, and Dr. Emmanuelle Genoyer for reagents and/or helpful discussion. We would like to thank Dr. Danielle Dauphars, Dr. Pam Fink, and all members of the Savan lab for manuscript review and editing. This work is funded by grants from the National Institutes for Health: K99AI175483 (N.S.G.); R01AI153246 (A.O.); R01AI155512 (S.M.H.); R01GM129090 (S-E.O.); R01AI145296, R21AI158788 (M.G.); R01AI175724, R21AI176442, and R21AI141823 (R.S.). Additionally, this work is funded by a Helen Hay Whitney Postdoctoral Fellowship (N.S.G.) and an American Cancer Society Postdoctoral Fellowship (M.G.T.). This work used an EASY-nLC1200 UHPLC and Thermo Scientific Orbitrap Fusion Lumos Tribrid mass spectrometer purchased with funding from a National Institutes of Health SIG grant S10OD021502 (S-E.O.).

## AUTHOR CONTRIBUTIONS

*Conceptualization:* N.S.G. and R.S.; *Investigation:* N.S.G., K.S., M.G.T., R.K.S., and D.M.M.; *Methodology:* N.S.G., K.S., M.G.T., S.D., M.P., L.H.C., D.M.M., and S-E.O.; *Formal Analysis:* N.S.G., K.S., M.G.T., and D.M.M.; *Software:* N.S.G., K.S., M.G.T., and D.M.M.; *Visualization:* N.S.G., K.S., and M.G.T.; *Resources:* A.O., S.M.H, S-E.O., M.G., and R.S.; *Writing – original draft:* N.S.G.; *Writing – review and editing:* N.S.G., K.S., M.G.T., R.K.S., D.M.M., L.H.C., S.D., A.O., S.M.H., S-E.O, M.G., R.S.; *Funding acquisition:* N.S.G., M.G.T., A.O., S.M.H., S-E.O., M.G., and R.S.

## SUPPLEMENTARY MATERIALS

Supplementary Figures S1-S6.

Supplementary Data S1 - APOBEC1-mediated profiling of MAVS associated RNAs

Supplementary Data S2 - IP-MS for RNA-modulated MAVS interactors

Supplementary Data S3 - List of primers, gene fragments, and siRNAs used in this study

## MATERIALS AND METHODS

### Cell lines

HEK293T (shortened to 293T throughout this manuscript) and NIH3T3 cells were grown in Dulbecco’s modification of Eagle’s medium (DMEM; Thermo Fisher) supplemented with 10% fetal bovine serum (HyClone), 25 mM HEPES (Thermo Fisher), 1X non-essential amino acids (Thermo Fisher), and 1X PSG (Thermo Fisher). Cells were continuously verified as mycoplasma free using the LookOut Mycoplasma PCR detection kit (Sigma-Aldrich).

### Plasmids and cloning

*MAVS and RLR pathway constructs*: pEFTak empty vector and FLAG-MAVS, and pEFBos FLAG-N-RIG have been described before (*55*). pEFTak FLAG-MAVS ΔCARD, FLAG-MAVS ΔTM, FLAG-MAVS ΔCARD+ΔTM and FLAG-MAVS Δ103-467 were generated by PCR mutagenesis of pEFTak FLAG-MAVS with linkers on primers for in-frame cloning. pEFTak FLAG-murineMAVS was cloned using InFusion cloning (Takara) of a PCR fragment of the MAVS open reading frame amplified from NIH3T3 cell cDNA and *NotI*- and *PmeI*-digested pEFTak FLAG vector. *APOBEC1 constructs:* pEFTak APOBEC1 was generated using InFusion cloning of the murine APOBEC1 open reading frame (gift of Dr. Jennifer Hyde) amplified by PCR, annealed primers coding for a 15 amino acid linker (protein sequence: SGSETPGTSESATPE), and *NotI*- and *PmeI*-digested pEFTak FLAG vector. Similarly, *NotI*- and *PmeI*-digested pEFTak FLAG vector, annealed primers for the linker, and PCR fragments of the entire MAVS coding sequence or the terminal 40 amino acids of MAVS were recombined using InFusion cloning to generate pEFTak APOBEC1-MAVS^FL^ and pEFTak APOBEC1-MAVS^C-term^ respectively. *3xFV-N-RIG:* pRRL 3xFV-N-RIG lentiviral vector was constructed by using InFusion cloning of a gene fragment (IDT) containing the N-terminal CARDs of murine RIG-I (amino acids 1-229) downstream of 3 tandem copies of FKBP12 carrying the F36V mutation (called “FV” domains) into *BamHI*-cut pRRL-Blast vector. 2 of the 3 copies contained silent mutations to prevent DNA recombination. *IFNB1*^2X^-GLuc: pTRIPZ *IFNB1*^2X^-GLuc reporter lentiviral vector was constructed by InFusion cloning of PCR amplified *IFNB1* promoter enhanceosome elements (−110 to −37 upstream of the transcription start site (Twist gene fragment)), ligated together with *BamHI*, into *SmaI*-cut pTRIPZ ISRE-GFP (Gift of Dr. Nicholas Heaton) (*53*). Further, the GFP gene was replaced with codon-optimized *Gaussia* luciferase open reading frame fused to the mODC-PEST domain (Twist gene fragment) by InFusion cloning into *XhoI-* and *XbaI*-cut vector. All primers used for cloning are provided in Supplementary Data S3.

### Generation of 293T RIG-I^KO^, 293T MAVS^KO^, and 293T IRF3^KO^ cells

293T cells deleted for RLR pathway proteins were generated by CRISPR/Cas9-mediated gene editing. RIG-I^KO^: Two guides targeting just upstream of the transcription start site (GCTAGTGAGGCACAGCCTGCGGG) and in exon 1 (CCCAGGTT-TGTGGTAAGATCTCC), respectively. MAVS^KO^: Single guide targeting exon 5 (CCTCAGCCCTCTGACCTCCAGCG). IRF3^KO^: Single guide targeting exon 2 (CCACTGGTGCATATGTTCCC) (*61*, *62*). 293T cells were transfected with the respective guide containing plasmids (pX330-sgRIGI-1 and pX330-sgRIGI-2, pX330-sgMAVS, or pX330-sgIRF3) along with pcDNA-Blast (encoding blasticidin resistance). Individual clones were selected following 0.2 μg/mL blasticidin selection and then screened for RIG-I, MAVS, or IRF3 protein expression by immunoblot. Clones with loss of protein expression were further validated by sequencing of amplicons spanning targeted regions.

### Generation and stimulation of 293T 3xFV-N-RIG cells

Lentivirus for 3xFV-N-RIG expression was produced in 293T cells by co-transfection of pRRL 3xFV-N-RIG, pMD2.G, and psPAX2. Stable 293T^WT^ and 293T IRF3^KO^ cell lines transduced with this lentivirus were selected using 0.5 µg/mL blasticidin. Cell lines were maintained in cDMEM supplemented with 0.2 µg/mL blasticidin. RLR signaling was activated by treatment with 10 nM B/B homodimerizer (AP20187; Clonetech) for the indicated amount of time.

### Generation of 293T *IFNB1*^2X^-GLuc cells

Lentivirus for *IFNB1*^2X^-GLuc expression was produced in 293T cells by co-transfection of pTRIPZ *IFNB1*^2X^-GLuc, pMD2.G, and psPAX2 and was used transduce 293T cells. Single transduced cells were cloned by dilution and colony outgrowth, and clones were screened for their ability to produce GLuc in the supernatant using the *Gaussia* Glow kit (Thermo Fisher).

### RT-qPCR

RNA was extracted using TRIzol (Thermo Fisher) or RNA columns (Machery-Nagel), and cDNA was generated using the PrimeScript RT-PCR kit (Takara). RT-qPCR was performed using a Viaa7 Real Time PCR instrument (Applied Biosciences) using TaqMan Universal PCR master mix II - UNG (Thermo Fisher). Primers for RT-qPCR are described in Supplementary Data S3.

### Immunoblotting

For sucrose gradient ultracentrifugation, immunoprecipitation, and MAVS activity assays, protein samples in assay-specific buffers were prepared with 1-2X final concentration of Laemmli buffer (Bio-Rad) with 2.5% β-ME and immunoblotted as described below. Whole cell lysates were prepared in a modified RIPA buffer (10 mM Tris pH 7.4, 150 mM NaCl, 0.5% sodium deoxycholate, and 1% Triton X-100) supplemented with protease-phosphatase inhibitor cocktail (Sigma-Aldrich) and clarified by centrifugation at 7500 xg for 10 mins at 4°C. Protein concentration was determined by Bradford assay (Bio-Rad). Equal amounts of protein sample in 1X Laemmli buffer with 2.5% β-ME were boiled and resolved by SDS-PAGE (Tris-Glycine gels; Bio-Rad) and transferred to methanol-activated PVDF membranes (Bio-Rad) by wet transfer. For probing of hard-to-detect targets, transferred membranes were fixed in membrane fixation buffer (7% acetic acid, 3% glycerol, 40% ethanol in water) prior to blocking with 3% BSA in tris-buffered saline with 0.1% Tween (TBS-T). Transferred membranes were incubated with relevant primary antibodies in 3% BSA in TBS-T with shaking for 1-2 hrs at room temperature or overnight at 4°C. After washing three times with TBS-T, membranes were incubated with species-specific horseradish peroxidase-conjugated secondary antibodies (Jackson, 1:5000). Chemiluminescence was detected using a Bio-Rad ChemiDoc XRS+ imaging instrument. The following primary antibodies were used for immunoblot: rabbit anti-MAVS (Bethyl and CST, 1:1000), mouse anti-MAVS (AdipoGen, 1:1000), rabbit anti-TRAF2 (CST, 1:1000), rabbit anti-TRAF3 (CST, 1:1000), mouse anti-TRAF6 (Santa Cruz, 1:250), rabbit anti-TRAF6 (CST, 1:1000), mouse anti-RIG-I (Novus, 1:1000), rabbit anti-CANX (CST, 1:1000), mouse anti-HuR (Santa Cruz, 1:500), rabbit anti-IRF3 (CST, 1:1000), rabbit anti-IRF3 phospho-S386 (Abcam, 1:1000), rabbit anti-FKBP12 (Pierce, 1:1000), rabbit anti-TOMM70 (Proteintech, 1:1000), rabbit anti-GAPDH (CST, 1:1000), rabbit anti-Tubulin (CST, 1:1000), HRP-conjugated anti-FLAG M2 (Sigma-Aldrich, 1:5000), and HRP-conjugated anti-β-actin (CST, 1:5000). Gels were quantified by densitometry using FIJI (*63*).

### Subcellular fractionation for crude mitochondrial extracts

For subcellular fractionation of crude mitochondrial extracts, 293T cells were harvested by pipetting in PBS, resuspended in 500 µL cold hypotonic buffer (10 mM Tris pH 7.4, 10 mM KCl, 1.5 mM MgCl_2_, 0.5 mM EGTA, and 1X protease-phosphatase inhibitor) for 15 mins on ice, and were then disrupted by 70 strokes of a tight pestle in 1mL Dounce homogenizers (Omni). Alternatively, cells were collected in cold PBS supplemented with 1X protease-phosphatase inhibitor for 15 mins on ice and were then disrupted by passing through a 27^1/2^-gauge syringe needle 10 times. These mechanically disrupted lysates were centrifuged at 1000 xg for 5 mins at 4°C to pellet nuclei and cell debris. Supernatants were then transferred to new tubes and centrifuged at 10000 xg for 10 mins at 4°C. Supernatants (cytosolic fraction; “cyto”) were transferred to new tubes or discarded. Pellets (crude mitochondrial fraction; “mito”) were resuspended in assay-specific buffers (see below). Protein concentration in cytosolic and crude mitochondrial fractions was measured by Bradford Assay.

### MAVS activity assay for *in vitro* phosphorylation of IRF3

MAVS activity assay was adapted from Zeng et al., using IRF3^KO^ cells to ensure that 293T^WT^ cytoplasmic lysates are the only source of IRF3 in the system in place of ^35^S-labeled IRF3 (*27*). The MAVS signalosome was activated in 293T IRF3^KO^ or 293T 3xFV-N-RIG IRF3^KO^ cells cultured in 10 cm plates by SenV infection (100 HAU/mL, 16 hpi) and B/B treatment (10 nM, 3 hpt), respectively. 293T^WT^ cells were concurrently cultured in a 10 cm plate and were left unstimulated. 5% of cells were stored for immunoblotting of whole cell lysates. Cells were fractionated as described above to generate cytosolic extract (“Cyto”) from 293T^WT^ cells and crude mitochondrial pellets from IRF3^KO^ cells. Mitochondrial pellets (“Mito”) were washed once with 200 µL mitochondrial reconstitution buffer (MRB; 20 mM HEPES-KOH pH 7.5, 10% glycerol, 0.5 mM EGTA), and resuspended in 50 µL MRB with 1% DDM and 1X protease-phosphatase inhibitor. After incubation on ice for 10 mins, mitochondrial and cytosolic lysates were centrifuged at 5000 xG at 4°C for 5 mins to pellet debris. 10 µg of mitochondrial lysate was mixed with RNaseIN as control, or with RNase, and were incubated for 15 mins on ice. 50 µg of cytosolic extract from 293T^WT^ cells was added to each sample in a final volume of 30 µL. 10 µL of 4X MAVS activity buffer (80 mM HEPES-KOH pH 7.0, 10 mM ATP, 20 mM MgCl_2_) was then added, and samples were incubated for 1 hr at 30°C with gentle shaking (250 rpm) in a Thermomixer. Following SDS-PAGE, transfer to PVDF membranes, and membrane fixation, samples were immunoblotted for IRF3 p-S386, IRF3, and MAVS.

### RNase-dependent shift assays by sucrose gradient ultracentrifugation

Sucrose gradient ultracentrifugation to identify RNase-dependent shift was performed as described by Caudron-Herger et al. with modifications (*31*). Sucrose gradients in 14 mL ultracentrifuge tubes (Beckman-Coulter) were prepared by sequential addition of 5-50% sucrose in 5% steps in gradient buffer (10 mM Tris pH 7.4, 100 mM NaCl, 1 mM EDTA), with freezing between the addition of each fraction (1.2 mL per fraction). Cytosolic and crude mitochondrial fractions from 293T, 293T RIG-I^KO^, or NIH3T3 cells in 10 cm plates with the indicated treatments (SenV infection, 100 HAU/mL, 14 hpi; transfected with pEFTak FLAG-MAVS or pEFTak FLAG-murineMAVS or pEFBos FLAG-N-RIG, 10 µg, 16 hpt using TransIT X2 (Mirus) transfection reagent) were isolated as described above. Mitochondrial pellets were lysed in PBS with 1% n-dodecyl-β-D-maltoside (DDM; Sigma-Aldrich) and 1X protease-phosphatase inhibitor, and 10% cytosolic extracts were added back to mitochondrial lysates. Following protein quantification, equal amounts (∼500 µg) of each sample were treated with RNaseIN as control (Promega, 40 U), or an RNase cocktail (per sample: RNase A (Thermo Fisher, 1 µg), RNase I (Thermo Fisher, 5 U), RNase T1 (Thermo Fisher, 500 U) and RNase III (Thermo Fisher, 1 U)) for 1 hr on ice. Treated samples were loaded on 5-50% sucrose gradient columns and ultracentrifuged using an SW40Ti rotor at 40000 xg for 18 hrs at 4°C. Following ultracentrifugation, equal fractions (1.25 mL) were transferred into new tubes by careful pipetting from the top of each tube. Protein in each fraction was precipitated by mixing 0.7 mL of each fraction with 1.5 mL of 100% ethanol and 0.5 µL GlycoBlue (Thermo Fisher) at −20°C overnight. Precipitated protein was pelleted by centrifugation at 13000 xg for 30 mins at 4°C, washed once with 70% ethanol, and dried. Protein pellets were resuspended in 1X Laemmli buffer with β-ME and dissolved by boiling at 95°C for 10 mins with shaking in a Thermomixer. Equal volume of each sample was resolved by SDS-PAGE and immunoblotted with the relevant antibodies.

### irCLIP to detect crosslinked RNA-protein complexes

To test the RNA-binding ability of MAVS and MAVS constructs, irCLIP was performed as described by Zarnegar et al. with slight modifications (*34*). For endogenous MAVS irCLIP, 293T^WT^ cells were cultured in 6-well plates and infected with SenV (100 HAU/mL, 16 hpi). For FLAG-tagged MAVS constructs (Fig. 2C, Supplementary Fig. S2), 293T MAVS^KO^ or NIH3T3 cells cultured in 6-well plates were transfected with the indicated FLAG-tagged plasmids for 16 hrs. For irCLIP of overexpressed MAVS constructs, mitochondrial extracts were prepared from 293T MAVS^KO^ cells as described above. At time of harvest, cells were washed with PBS, crosslinked with 254 nm UV-C light (0.15 J/cm^2^), and lysed in 200 µL irCLIP lysis buffer (50 mM Tris pH 7.5, 150 mM NaCl, 1 mM EDTA, 1% Triton X-100, 0.1% SDS, 1X protease inhibitor). After sonication in ice slurry, lysates were clarified by centrifugation at 5000 xg for 10 mins at 4°C, quantified by Bradford assay, and concentrations were equalized. 5% of each sample was stored for input control. 200 µg of each lysate was incubated with 20 µL Protein G Dynabeads (Thermo Fisher) pre-conjugated with 4 µg mouse anti-FLAG M2 antibody (Sigma-Aldrich) or rabbit anti-MAVS (Bethyl) in a final volume of 500 µL irCLIP lysis buffer for 2 hrs at 4°C with rotation. Beads were then sequentially washed with the following ice-cold buffers: twice with 1 mL irCLIP lysis buffer, twice with 1 mL high stringency buffer (20 mM Tris pH 7.5, 120 mM NaCl, 25 mM KCl, 5 mM EDTA, 1% Triton X-100, 1% sodium deoxycholate, 0.1% SDS), twice with 1 mL high salt buffer (20 mM Tris pH 7.5, 500 mM NaCl, 5 mM EDTA, 1% Triton X-100, 1% NaDOC), twice with with 1 mL low salt buffer (20 mM Tris pH 7.5, 5 mM NaCl, 5 mM EDTA, 1% Triton X-100), and twice with 1 mL NT2 buffer (50 mM Tris pH 7.5, 150 mM NaCl, 1 mM MgCl_2_, 0.05% NP-40). 10% beads were then removed for immunoblot analysis, and the rest of the beads were then resuspended in 30 µL NT2 buffer containing 25 ng/mL RNase A and 15% PEG400 (Sigma-Aldrich) for on-bead RNase digestion at 30°C for 15 mins with shaking (1200 rpm) in a Thermomixer. RNase digestion was quenched by the addition of 1 mL high stringency buffer. Beads were washed twice with 0.3 mL PNK wash buffer (50 mM Tris pH 7.0, 10 mM MgCl_2_), and then resuspended in 30 µL PNK dephosphorylation mix (1X PNK buffer (Promega), 0.5 µL RNaseIN, 1 µL T4 PNK (Promega), 4 µL PEG400). Dephosphorylation reactions were conducted at 37°C for 60 mins with shaking (1200 rpm) in a Thermomixer. Dephosphorylation mix was removed, and beads were washed with 0.3 mL PNK wash buffer. For ligation of IRDye-800 conjugated oligo to RNA crosslinked to protein, beads were resuspended in 30 µL RNA ligation mix (1X RNA ligase I buffer (NEB), 1 µL RNA ligase I (NEB), 0.5 µL IRDye800-labeled oligonucleotide (*34*), 5 µL PEG400, and 0.5 µL RNaseIN) and incubated for 16 hrs at 16°C with shaking in Thermomixer (1200 rpm). Ligation mix was then removed, and beads were washed twice with 0.3 mL PNK wash buffer, prior to elution of RNA-protein complexes in 20 µL 1X LDS Buffer (Thermo Fisher) + 10% β-ME at 80°C for 10 mins. 5 µL of eluates, as well as input controls, were then resolved by SDS-PAGE on 4-12% Bis-Tris NuPAGE gels (Thermo Fisher) and transferred to nitrocellulose membranes. Fluorescent RNA-protein complexes in the eluates were visualized on a LiCOR Odyssey FC imager. The same membranes were then subjected to immunoblotting using HRP-conjugated anti-FLAG M2 or anti-MAVS antibody.

### Prediction of disorder conservation in MAVS

Prediction of intrinsic disorder in human MAVS was conducted using IUPred3 (*36*) and FuzPred (*64*) web portals. The IUPred3 package was used for the analysis of disorder conservation in MAVS homologs downloaded from NCBI and Ensembl.

### Identification of MAVS-associated RNAs using APOBEC1-mediated profiling

293T cells seeded in 12-well plates were transfected with 0.5 µg plasmid encoding APOBEC1, APOBEC1-MAVS^C-term^, and APOBEC1-MAVS^FL^ using the Mirus X2 transfection reagent in biological triplicate. At 24 hrs post-transfection, RNA was extracted using TRIzol, treated with Turbo DNase I (Thermo Fisher), cleaned by phenol-chloroform extraction, and precipitated. Total RNA stranded library preparation with rRNA depletion and paired end 100 bp sequencing using the DNBseq platform was performed through BGI Genomics.

RNA-sequencing data was subjected to quality control and adaptor trimming using SOAPnuke (*65*). Additional quality control was performed using fastqc and multiqc (*66*, *67*); all samples passed thresholding and were retained. A snakemake pipeline using samtools, Rsamtools, bedtools, and pybedtools was used was used to orchestrate analysis (https://github.com/ksomf/seq_pipeline) (*68–72*). Reads were aligned to GRCh38 using STAR (*73*). The Bullseye pipeline (https://github.com/mflamand/Bullseye) was used to detect C-to-T editing at sites with at least 10 reads of coverage per sample, with an edit ratio between 2% and 95% (minEdit=5), and an edit ratio at least 1.5-fold higher than control samples (editFoldThreshold=1.5) (*46*, *47*).

### Validation of *PMAIP1* editing by Sanger sequencing

293T cells were transfected with 0.5 µg plasmids encoding APOBEC1-MAVS^C-term^ and APOBEC1-MAVS^FL^ for 24 hrs using Mirus X2. After RNA extraction and cDNA preparation using the PrimeScript RT kit, the *PMAIP1* 3′UTR was amplified by PCR (Primers in Supplementary Data S3), and purified amplicons were subjected to Sanger sequencing. The MultiEditR web tool was used to analyze C-to-T editing in .ab1 chromatograms at sites called by Bullseye (*74*).

### CLIP-RT-qPCR

293T cells seeded in 6-well plates were transfected with plasmids encoding FLAG-MAVS or FLAG-MAVS Δ103-467 using Mirus TransIT X2. At 24 hpi, cells were washed with PBS, crosslinked with 254 nm UV-C light (0.15 J/cm^2^), and lysed in 200 µL irCLIP lysis buffer supplemented with protease-phosphatase and RNase inhibitors. After sonication in ice slurry, lysates were clarified by centrifugation at 5000 xg for 10 mins at 4°C, quantified by Bradford assay, and concentrations were equalized. After removing 20 µg lysate for input RNA (10%), 200 µg lysates were incubated with washed Protein G Dynabeads bound to anti-FLAG antibody (25 µL beads and 4 µg antibody per sample) in irCLIP lysis buffer for 2 hrs at 4°C with rotation. Beads were then washed twice with 1 mL irCLIP lysis buffer, twice with 1 mL high stringency buffer, and twice with PBS. Beads were resuspended in 100 µL PBS, and 10% beads were removed for immunoblot analysis. After the addition of 10 µL 10X Turbo DNase buffer and 2 µL Turbo DNase I, beads were incubated at 37°C with shaking (1000 rpm) in a Thermomixer. RNA was then eluted from beads by incubation with 5 µg Proteinase K (Thermo Fisher) at 50°C with shaking (1000 rpm). After removal of supernatant, beads were washed twice with 100 µL PBS and washes were pooled with the eluates. RNA was purified from input and IP samples using TRIzol LS (Thermo Fisher). Equal volumes of eluted RNA were used for cDNA synthesis, quantified by RT-qPCR, and normalized to RNA levels in input samples. CLIP-RT-qPCR of RNA bound to endogenous MAVS was performed similarly from uninfected and SenV-infected (100 HAU/mL, 20 hours) 293T cells, except that lysates were incubated with Protein G Dynabeads bound to rabbit anti-MAVS or normal rabbit IgG. Fold enrichment was calculated with the enrichment over input in FLAG-MAVS Δ103-467 for each IgG set as 1.

### Immunofluorescence microscopy

293T cells seeded in 8-well chamber slides (Millipore) were transfected with the indicated FLAG-tagged APOBEC1-MAVS or control constructs. At 24 hrs post-transfection, cells were washed with PBS and fixed with 4% paraformaldehyde for 10 mins. Cells were permeabilized with 0.2% Triton X-100 in blocking buffer (3% BSA in PBS). Permeabilized cells were incubated with primary anti-FLAG antibody (1:2500) for 1 hr, washed with PBS, and were then incubated in AlexaFluor conjugated secondary antibody (1:2500; Thermo Fisher) and DAPI (1:1000) for 1 hr. After further PBS washes, chamber slides were mounted using ProLong Gold (Thermo Fisher). Images were acquired using a Nikon C2 confocal microscope.

### SDD-AGE for MAVS oligomerization

Crude mitochondrial pellets were generated from SenV-infected (200 HAU/mL, 12 hpi), FLAG-MAVS-transfected (10 µg, 16 hpi, TransIT X2 transfection reagent), or unstimulated 293T cells as described above. Mitochondrial pellets were resuspended in 50 µL PBS, divided equally into three separate tubes, and treated with RNaseIN, RNase cocktail, or 30 µM β-ME. Following incubation on ice for 30 mins, an equal volume of 2X SDD-AGE sample buffer (2% SDS, 10% glycerol, 0.02% bromophenol blue in 0.5X tris-borate-EDTA (TBE)) was added to each tube. 10% of sample was stored for input controls, supplemented with Laemmli buffer, boiled, and immunoblotted by SDS-PAGE. 50% of the sample was loaded on a 1.5% agarose gel prepared in 1X TBE buffer. Samples were resolved by vertical electrophoresis (70V, 120 mA, 1 hr) in SDD-AGE running buffer (1X TBE, 0.1% SDS), and transferred (20V, 250 mA, constant voltage, overnight at 4°C) to PVDF membranes in SDD-AGE transfer buffer (25 mM Tris (pH 8.3), 192 mM glycine, 20% (v/v) ethanol). Transferred membranes were fixed with 0.25% glutaraldehyde in PBS for 20 mins with shaking and immunoblotted with anti-FLAG or anti-MAVS primary antibodies.

### Immunoprecipitation and mass spectrometry

293T cells were seeded in two 10 cm plates per condition and transfected with pEFTak FLAG-MAVS or empty vector using the Mirus TransIT X2 transfection reagent. Cells were harvested at 20 hrs post-transfection and crude mitochondrial extracts were prepared by mechanical disruption using a 27^1/2^ gauge needle as described above. Crude mitochondrial pellets were lysed in 300 µL IP-MS lysis buffer (25 mM Tris pH 7.5, 150 mM NaCl, 5 mM MgCl_2_, 0.5% DDM) supplemented with protease-phosphatase inhibitor and RNase inhibitor. 0.5 mg of mitochondrial lysates were incubated with washed Protein G Dynabeads pre-bound to anti-FLAG antibody (35 µL beads with 8 µg antibody per sample) in IP-MS wash buffer (25 mM Tris pH 7.5, 150 mM NaCl, 5 mM MgCl_2_, 0.1% DDM) in a final volume of 1 mL for 4 hrs at 4°C with rotation. Beads were then washed twice with 1 mL IP-MS wash buffer. MAVS + RNase samples were then incubated with 0.5 mL IP-MS wash buffer containing RNase A (1 µg/mL) and RNase T1 (1 U/µL) for 30 mins at room temperature with rotation. Concurrently, empty vector and MAVS - RNase samples were incubated in 0.5 mL IP-MS wash buffer containing RNase inhibitor. Beads were then washed twice with 1 mL IP-MS wash buffer, and twice in wash buffer without DDM. After moving beads to fresh tubes, samples were subjected to on-bead reduction, alkylation, and tryptic digest. Bound proteins were reduced with 5 mM DTT (Sigma-Aldrich) in 50 mM TEAB (Thermo Fisher) in a final volume of 100 µL for 30 mins at room temperature, followed by alkylation with 10 mM iodoacetamide (Thermo Fisher) for 30 mins at room temperature in the dark. Proteins were digested with 1 µg MS-grade trypsin (Thermo Fisher) at 37°C overnight with gentle shaking (800 rpm) in a Thermomixer. After addition of another 1 µg trypsin, samples were similarly incubated for 4 hrs. After collecting post-trypsin supernatants, beads were washed twice with 50 µL of 50 mM TEAB and washes were pooled with digested peptides.

Digested peptide samples were acidified with trifluoroacetic acid (TFA) to a final concentration of 0.5% and then desalted using C18 StageTips according to the published protocol with minor adjustments (*75*). StageTips were activated with 50 µL methanol, followed by sequential addition of 50µL StageTip buffer B (80% acetonitrile (ACN) in 0.1% TFA) and 50 µL StageTip buffer A (0.1% TFA). After sample loading, peptides were washed with 50 µL Stage Tip buffer A. Peptides from StageTips were eluted using 50 µL 45% ACN, 0.1% TFA into a 96-well plate. Samples were then dried down by vacuum centrifugation and resuspended in StageTip buffer A. Peptides were separated on an EASY-nLC 1200 System (Thermo Fisher) using 20 cm long fused silica capillary columns (100 μm ID, laser-pulled in-house using the Sutter P-2000) packed with 3 μm 120 Å reversed phase C18 beads (Dr. Maisch). The liquid chromatography (LC) gradient was 90 mins long with 5−35% B at 300 nL/min. LC solvent A was 0.5% acetic acid and LC solvent B was 0.5% acetic acid, 80% ACN. MS data was collected with a Thermo Fisher Orbitrap Fusion Lumos using a data-independent acquisition (DIA) method with a 120K resolution Orbitrap MS1 scan and 12 m/z isolation window, and 30K resolution Orbitrap MS2 scans for precursors from 400-1000 m/z.

Data .raw files were converted to .mzML using MSConvert (v3.0.21251-d2724a5) and spectral libraries were built using MSFragger-DIA with FragPipe (v20.0) (*76*) with quantification through DIA-NN v1.8.2 (*77*). Database search was against the UniProt human database (updated July 6^th^, 2023) with supplemental spike-in of common contaminants, containing 20477 sequences and 20477 reverse-sequence decoys. For the MSFragger analysis, both precursor and initial fragment mass tolerances were set to 20 ppm and spectrum deisotoping, mass calibration, and parameter optimization were enabled. Enzyme specificity was set to “stricttrypsin” and up to two missed trypsin cleavages were allowed. Methionine oxidation, N-terminal acetylation, serine, threonine, and tyrosine phosphorylation, −18.0106 Da on N-terminal glutamic acid, and −17.0265 Da on N-terminal glutamine and cysteine were set as variable modifications. Cysteine carbamidomethylation was set as a fixed modification. The maximum number of variable modifications per peptide was set to 3. FragPipe/DIA-NN output files were processed and analyzed using the Perseus software package v2.0.7.0 (*78*). Rows were retained only if at least one of the three experimental groups had at least 3 of 5 valid values. Data imputation was performed using a modeled distribution of MS intensity values downshifted by 1.8 and having a width of 0.2. Expression values (protein MS intensities) were log_2_-transformed and normalized by subtracting the median log_2_ expression value from each expression value within each MS run. For statistical testing of significant differences in enrichment, a two-sample Student’s t test was applied.

Enriched proteins were filtered for common contaminants of IP-MS experiments using the CRAPome database, and only proteins found in < 30% of CRAPome datasets were chosen for further analysis (*79*). Interactome network analysis was performed and visualized using the STRING app in Cytoscape (*80*, *81*). Gene set enrichment analysis of MAVS-interacting proteins was performed on a pre-ranked list of identified proteins, with log_2_FC(MAVS_CTRL-EV) x −log_10_(p-value) as the ranking metric (*51*, *52*). Network analysis of enriched GO: Biological Process pathways was performed and visualized with Enrichment Map and Cytoscape (*81*, *82*).

### Targeted siRNA screening

293T *IFNB1*^2X^-GLuc cells plated in 96-well plates were transfected with 1 pmol/well siRNA (Silencer Select, Thermo Fisher) using Lipofectamine RNAiMAX (Thermo Fisher) in technical triplicate. At 36 hours post-siRNA transfection, media was exchanged, and cells were transfected with 50 ng/well (293T) or 20 ng/well (Huh7) of *in vitro* transcribed poly-U/UC RNA using the Mirus X2 reagent (*55*). At 24 hrs post-poly-U/UC RNA transfection, 10 µL supernatant from each well was transferred to white opaque 96-well plates, wherein it was mixed with *Gaussia* Glow assay buffer with 1X coelenterazine (Thermo Fisher), and luminescence was read on a Biotek Synergy plate reader. Cellular viability was measured using Cell Titer-Glo (Promega). Secondary screen for *IFNB1* mRNA induction was performed similarly in 293T cells seeded in 48-well plates. RNA was extracted at 24 hrs post-poly-U/UC RNA transfection using TRIzol and RT-qPCR was performed as described. All siRNA sequences or assay IDs are listed in Supplementary Data S3.

### Data availability

Raw APOBEC1-mediated profiling data have been submitted and are available through GEO (GSE244000). Raw mass spectrometry data have been deposited to and are available through the MassIVE repository (MSV000092912).

## SUPPLEMENTARY FIGURES

**Supplementary Fig. S1.**
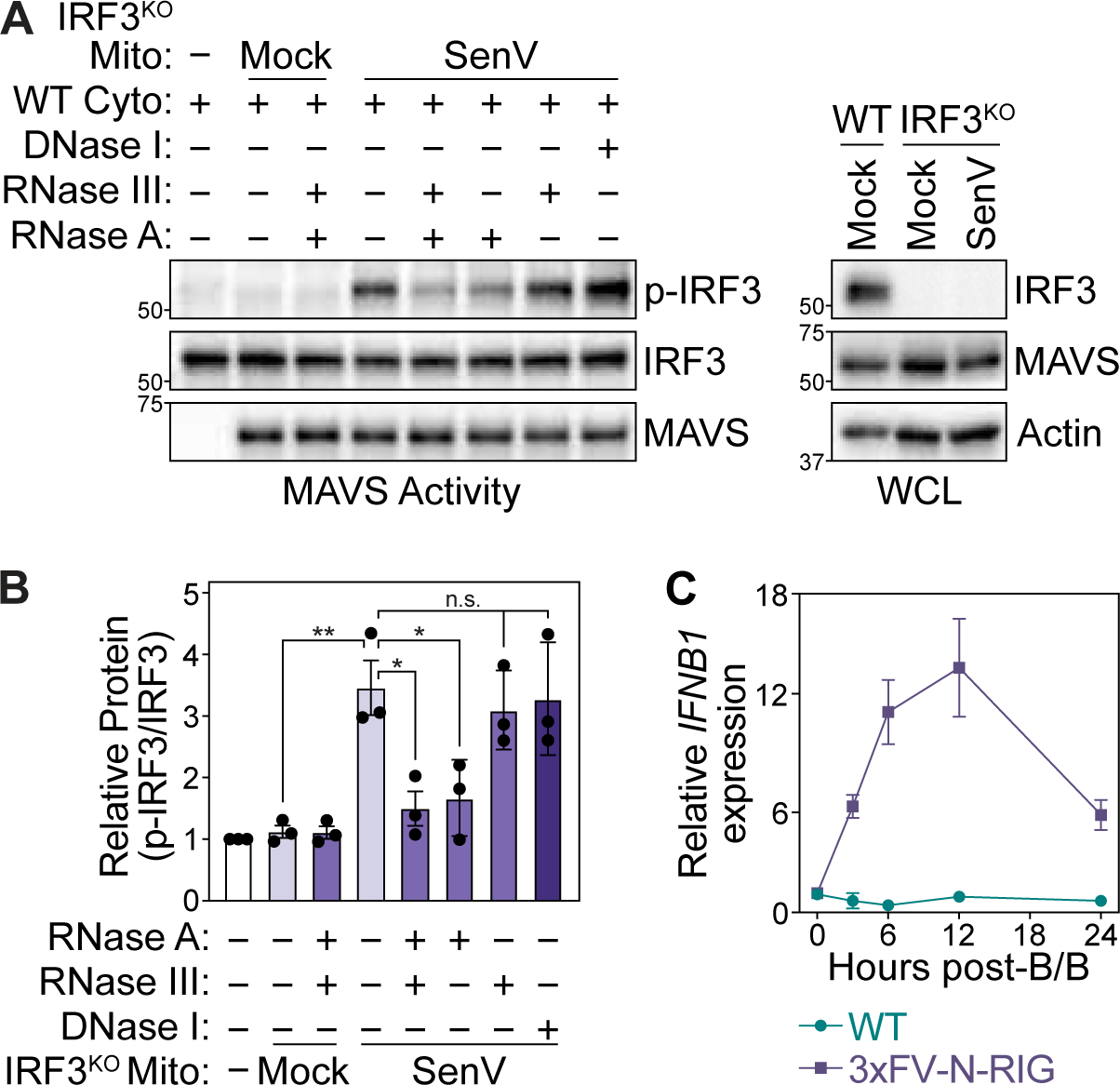
**(A)** MAVS activity assay to analyze *in vitro* IRF3 phosphorylation by mitochondrial extracts treated with the indicated nucleases from mock or SenV-infected (100 HAU/mL, 16 hpi) 293T IRF3^KO^ cells. Data are representative of 3 biological replicates. **(B)** Quantification of p-IRF3 (S386) relative to IRF3 from experiments in (A). Values are the ± SEM of 3 biological replicates. * p ≤ 0.05, ** p ≤ 0.01, *** p ≤ 0.001 by one-way ANOVA with Tukey’s multiple comparison test. **(C)** RT-qPCR for the expression of *IFNB1* RNA relative to *HPRT1* in 293T^WT^ cells and 293T cells stably expressing 3xFV-N-RIG treated with B/B (10 nM) over the indicated time course. Data are from a single biological replicate.

**Supplementary Fig. S2.**
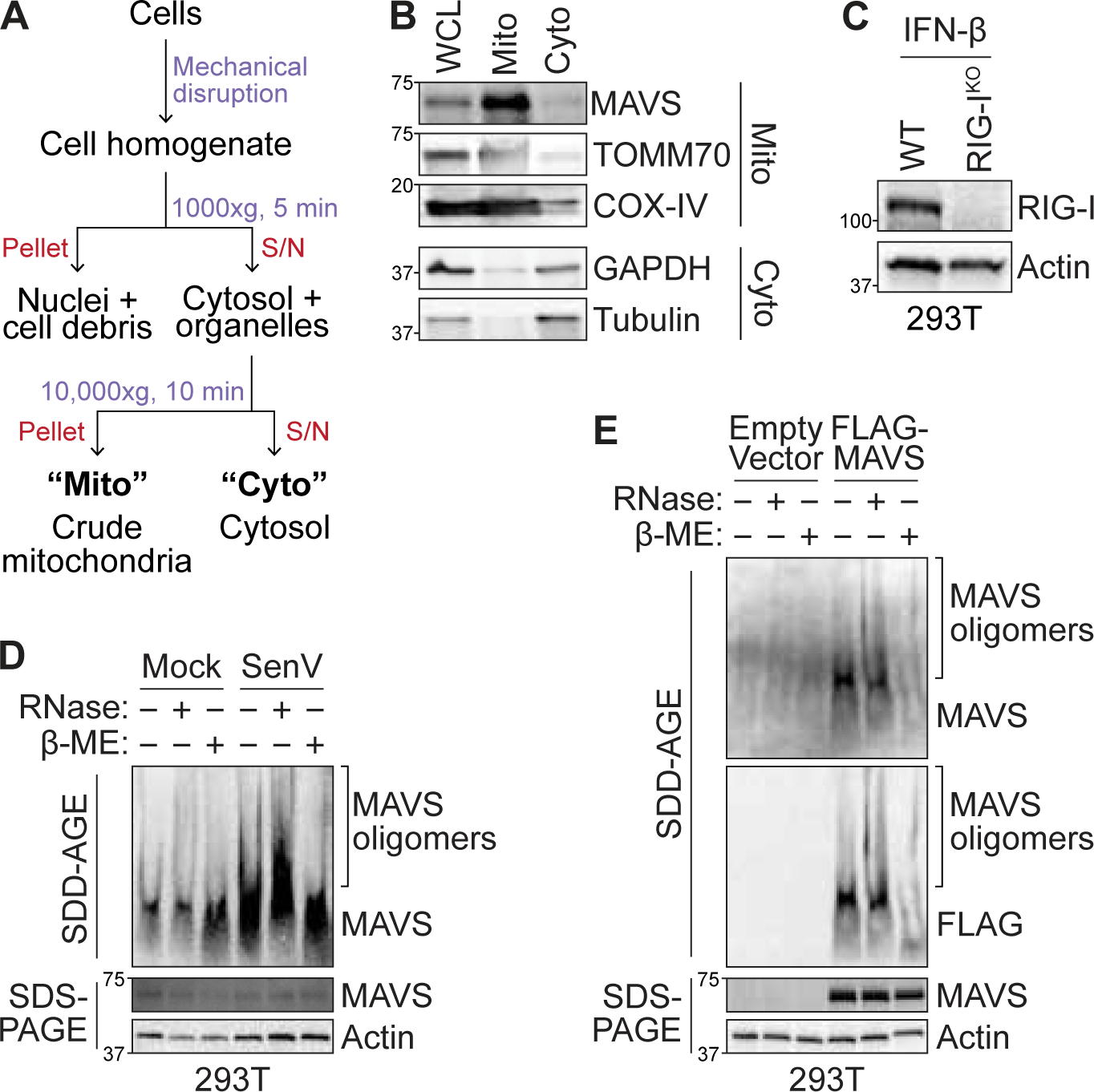
**(A)** Differential centrifugation strategy to isolate crude mitochondrial fractions. **(B)** Immunoblotting of whole cell (WCL), crude mitochondrial (Mito), and cytosolic (Cyto) fractions for mitochondrial and cytosolic proteins. Data are representative of 2 biological replicates. **(C)** Immunoblot of RIG-I expression in 293T^WT^ and 293T RIG-I^KO^ cells. Cells were treated with 100 U/mL IFN-β for 16 hrs to induce RIG-I expression. **(D)** SDD-AGE (for MAVS oligomers) and SDS-PAGE analysis of untreated, RNase-treated, or β-ME-treated mitochondrial extracts from mock- and SenV-infected (200 HAU/mL, 12 hpt) 293T cells. **(E)** SDD-AGE and SDS-PAGE analysis of untreated, RNase-treated, or β-ME-treated mitochondrial extracts from 293T cells transfected with empty vector or FLAG-MAVS (24 hpt). Data are representative of 2 ((B) and (C)) and 3 ((D) and (E)) biological replicates.

**Supplementary Fig. S3.**
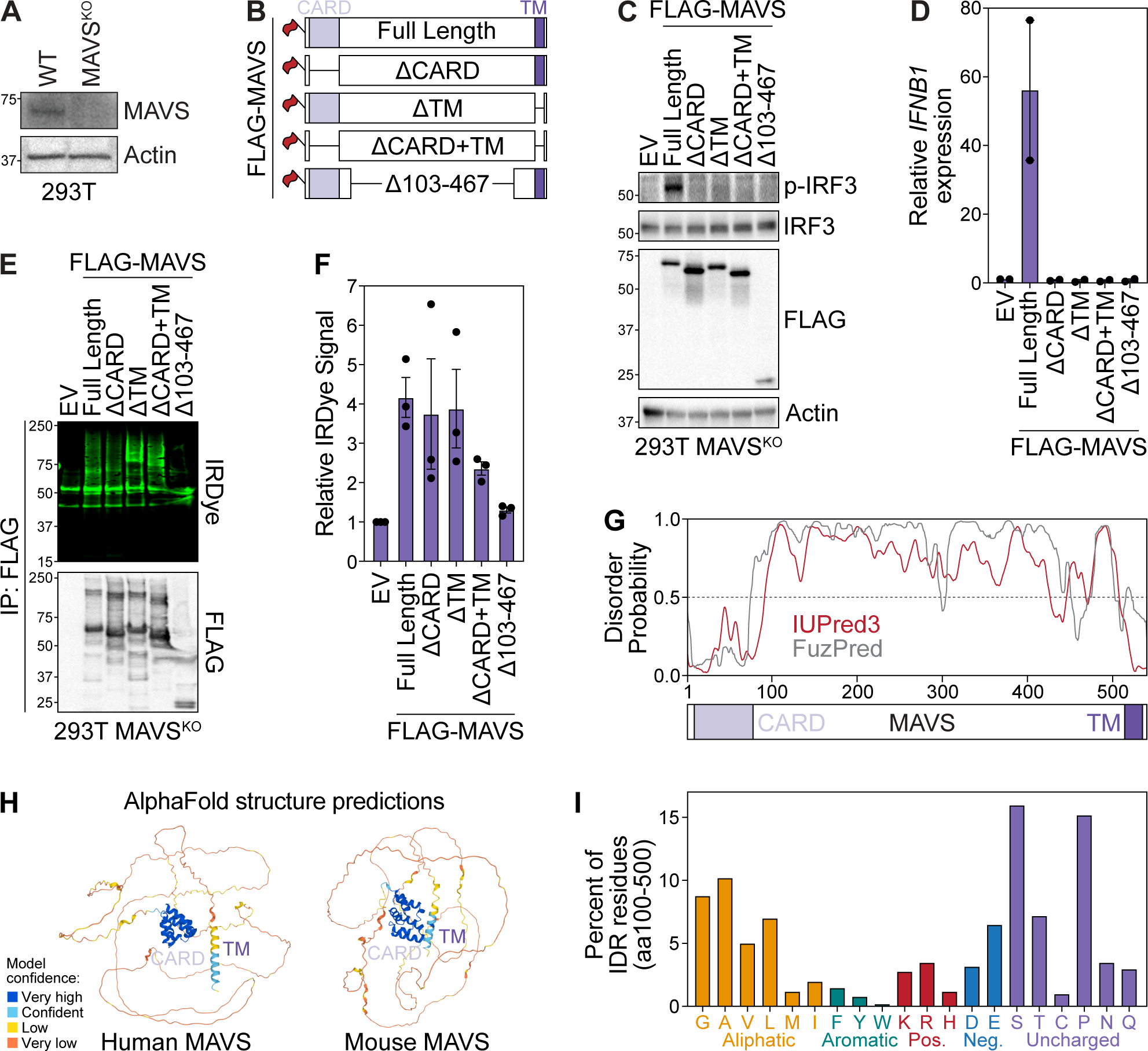
**(A)** Immunoblot of MAVS expression in 293T^WT^ and 293T MAVS^KO^ cells. **(B)** Schematic of MAVS constructs used in experiments in (C-F). **(C)** Immunoblot analysis of the indicated FLAG-tagged MAVS constructs expressed in 293T MAVS^KO^ cells (24 hpt). Data are representative of 2 biological replicates. **(D)** RT-qPCR analysis of *IFNB1* induction relative to *HPRT1* following expression of the indicated MAVS constructs in 293T MAVS^KO^ cells (24 hpt). Values are the mean ± SEM of 2 biological replicates. **(E)** irCLIP of the indicated FLAG-tagged MAVS constructs expressed in 293T MAVS^KO^ cells (24 hpt). Data are representative of 3 biological replicates. **(F)** Quantification of IRDye signal in experiments in (C). Values are mean ± SEM of 3 biological replicates. **(G)** Comparison of disorder predicted in human MAVS by IUPred3 (red) and FuzPred (grey). **(H)** AlphaFold structure predictions of human and murine MAVS. **(I)** Percent of each amino acid encoded in the MAVS disordered region (aa 100-500).

**Supplementary Fig. S4.**
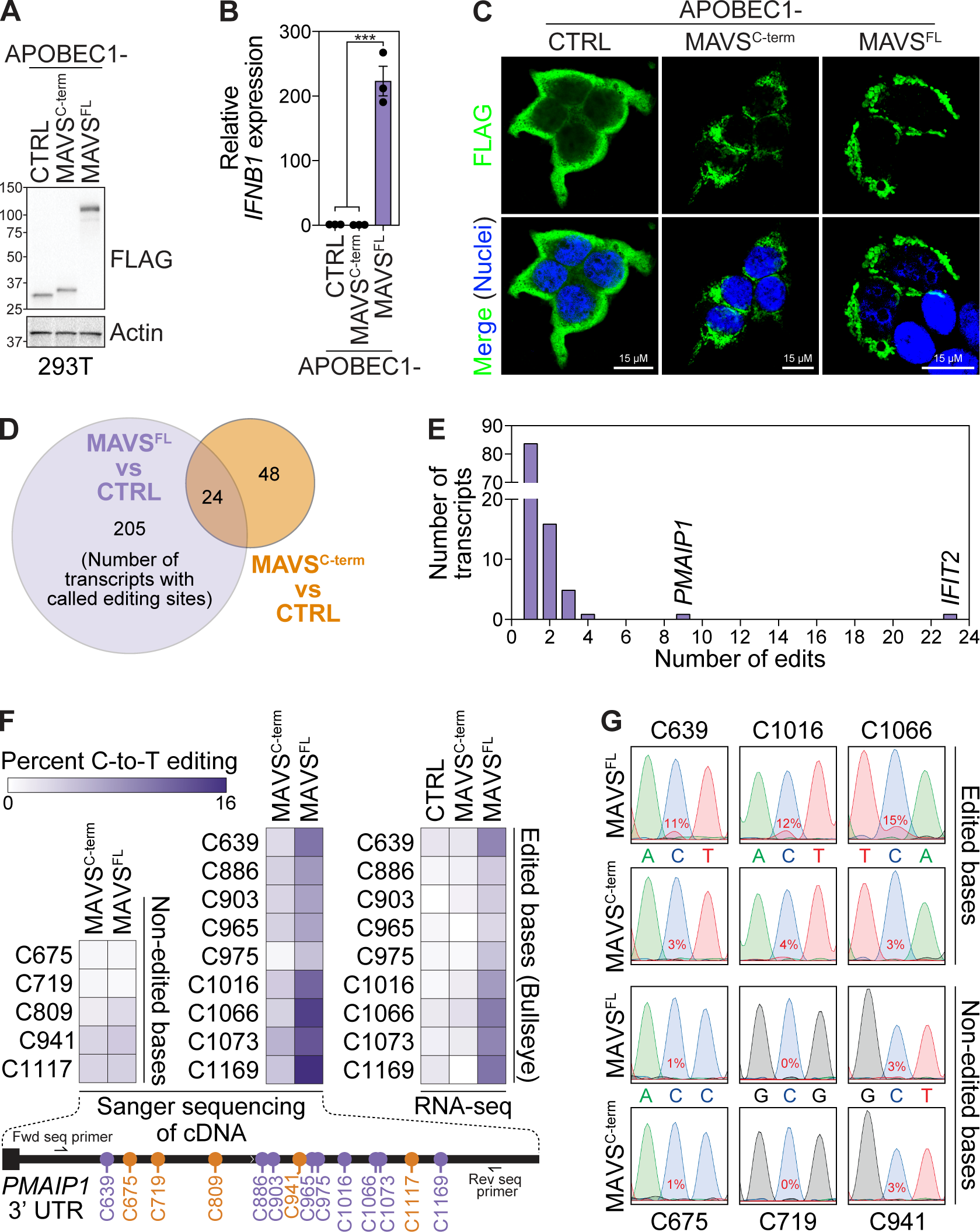
**(A)** Immunoblot analysis of the expression of transfected FLAG-APOBEC1 fusion constructs expressed in 293T cells (24 hpt). Data are representative of 3 biological replicates, from the same samples subjected to RNA-seq. **(B)** RT-qPCR analysis of *IFNB1* expression relative to *HPRT1* in 293T cells expressing FLAG-APOBEC1 fusion constructs. Values are mean ± SEM of 3 biological replicates. **(C)** Immunofluorescence micrographs of 293T cells transfected with FLAG-APOBEC1 fusion constructs (24 hpt) and stained for FLAG (green). Nuclei were stained with DAPI. **(D)** Venn diagram of the number of transcripts with called editing sites when comparing APOBEC1-MAVS^FL^ or APOBEC1-MAVS^C-term^ with APOBEC1 (CTRL). **(E)** Histogram of the number of transcripts with the indicated number of called editing sites when comparing APOBEC1-MAVS^FL^ with APOBEC1-MAVS^C-term^ + CTRL. **(F)** Sanger sequencing validation of 9 edited and 5 non-edited bases in the 3′ UTR of *PMAIP1*. **(G)** Sanger chromatograms of 3 representative edited and non-edited sites in the 3′ UTR of *PMAIP1*.

**Supplementary Fig. S5.**
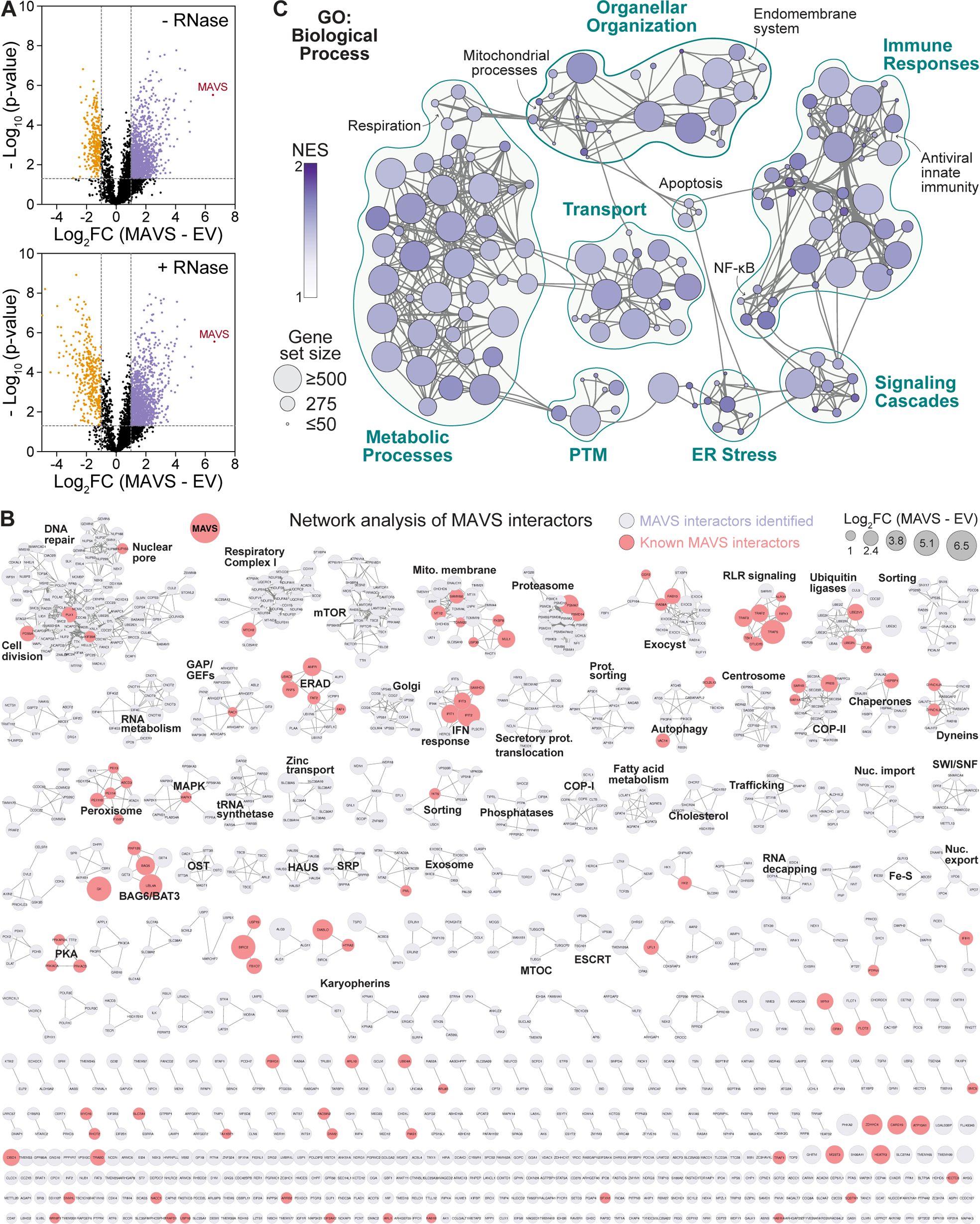
**(A)** Volcano plots of proteins enriched with MAVS −/+ RNase relative to empty vector controls by IP-MS. Purple and yellow dots represent significantly enriched or de-enriched (|Log_2_FC| ≥ 2, adjusted p ≤ 0.05) proteins respectively. **(B)** STRING functional network analysis of 1094 proteins significantly enriched with MAVS by IP-MS, and which are not common contaminants in the CRAPome database (identified in < 30% of CRAPome datasets). Clusters indicate proteins within known complexes or with shared function. Size of circle indicates Log_2_FC enrichment with MAVS. Proteins highlighted in red are known interactors of MAVS (based on the BioGRID database and literature search). **(C)** Network representation of Gene Set Enrichment Analysis results. Gene Ontology (GO): Biological Process terms related to immune responses, signaling cascades, intracellular transport, metabolic processes, organellar organization, ER stress, and post-translational modifications are enriched (FDR < 10%).

**Supplementary Fig. S6.**
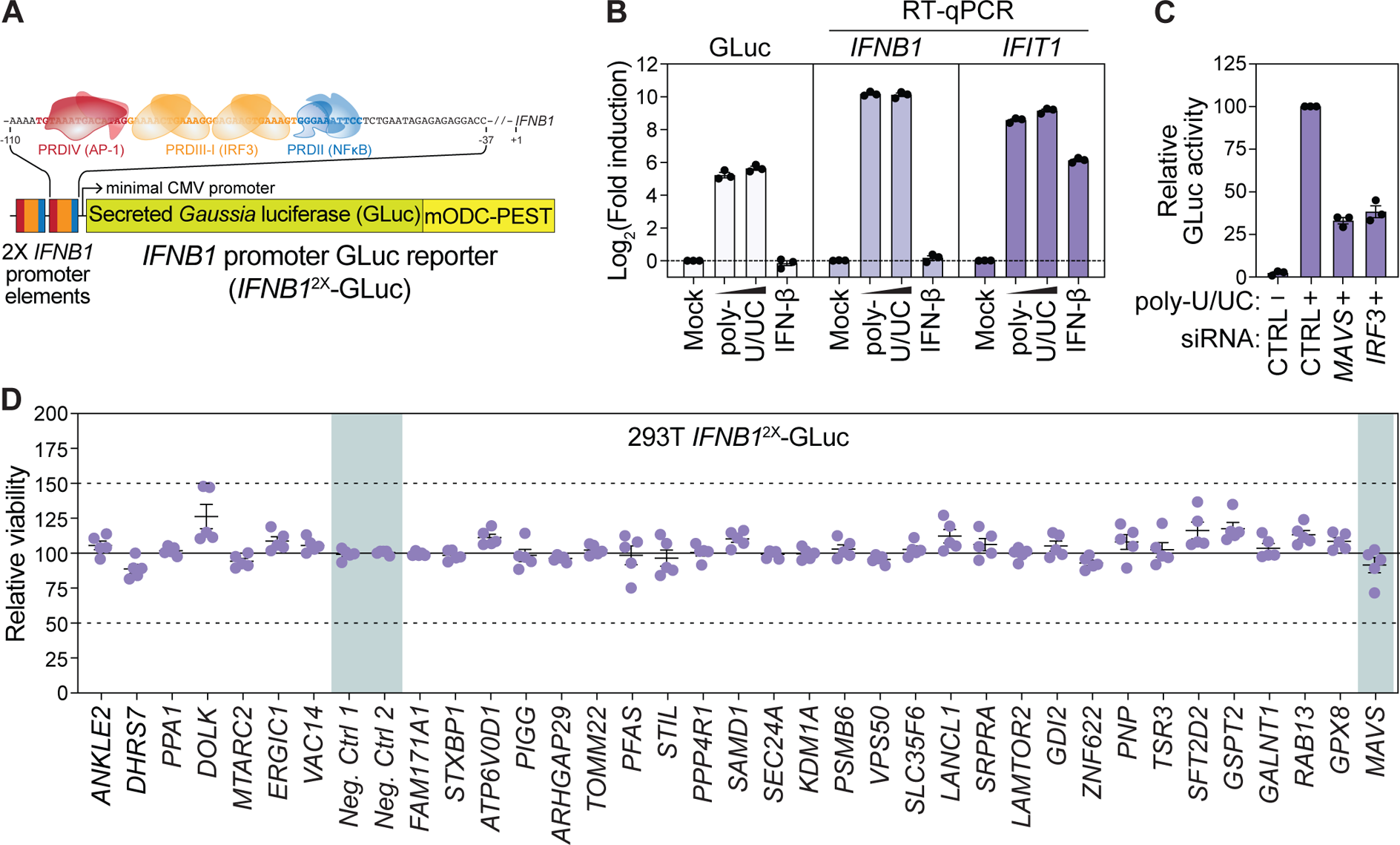
**(A)** Schematic of *IFNB1*^2X^-GLuc reporter used for functional screening. Two *IFNB1* promoter enhanceosome elements (nt 110 - 37 upstream of the transcription start site of *IFNB1*) containing binding sites for AP-1, IRF, and NF-κB transcription factors drive induction of a secreted *Gaussia* luciferase (GLuc) gene. A destabilizing PEST domain fused to GLuc reduces background expression. **(B)** Induction of GLuc or control transcripts *IFNB1* and *IFIT1* relative to *HPRT1* (measured by RT-qPCR) in clonal 293T cells stably expressing *IFNB1*^2X^-GLuc reporter with increasing amounts of transfected poly-U/UC RNA (0.5 µg and 1 µg) or IFN-β (1000 U/mL) for 24 hrs. **(C)** Induction of GLuc in clonal 293T cells stably expressing the *IFNB1*^2X^-GLuc reporter by transfected poly-U/UC RNA (0.5 µg, 24 hpt) in cells treated with the indicated siRNAs (36 hrs). **(D)** Viability as measured by Cell-Titer Glo of 293T *IFNB1*^2X^-GLuc cells transfected with the indicated siRNAs (36 hrs) and poly-U/UC RNA (50 ng, 24 hrs). Viability data were used to normalize data presented in Fig. 5E. Values are the mean ± SEM of 3 ((B) and (C)) or 5 (D) biological replicates respectively.

